# Engineered reactivity of a bacterial E1-like enzyme enables ATP-driven modification of protein C termini

**DOI:** 10.1101/2024.05.13.593989

**Authors:** Clara L. Frazier, Debashrito Deb, Amy M. Weeks

## Abstract

In biological systems, ATP provides an energetic driving force for peptide bond formation, but protein chemists lack tools that emulate this strategy. Inspired by the eukaryotic ubiquitination cascade, we developed an ATP-driven platform for C-terminal activation and peptide ligation based on *E. coli* MccB, a bacterial ancestor of ubiquitin-activating (E1) enzymes that natively catalyzes C-terminal phosphoramidate bond formation. We show that MccB can act on non-native substrates to generate an *O*-AMPylated electrophile that can react with exogenous nucleophiles to form diverse C-terminal functional groups including thioesters, a versatile class of biological intermediates that have been exploited for protein semisynthesis. To direct this activity towards specific proteins of interest, we developed the Thioesterification C-terminal Handle (TeCH)-tag, a sequence that enables high-yield, ATP-driven protein bioconjugation via a thioester intermediate. By mining the natural diversity of the MccB family, we developed two additional MccB/TeCH-tag pairs that are mutually orthogonal to each other and to the *E. coli* system, facilitating the synthesis of more complex bioconjugates. Our method mimics the chemical logic of peptide bond synthesis that is widespread in biology for high-yield *in vitro* manipulation of protein structure with molecular precision.

## Introduction

In living systems, ATP provides an energetic driving force for peptide bond synthesis. For example, activation of the α-carboxylate of amino acids or protein/peptide C termini by adenylation enables formation of ester and thioester intermediates that function in diverse pathways including ribosomal protein synthesis^1^, non-ribosomal peptide synthesis^2^, and the ubiquitination cascade^3^. Cleavage of ATP to AMP and pyrophosphate (PPi) provides a large thermodynamic driving force (ΔG°′ = −45.6 kJ/mol for hydrolysis) for otherwise unfavorable biosynthetic reactions^4^. In both protein translation and non-ribosomal peptide synthesis, amino acids are initially activated by ATP-dependent adenylation to hydrolytically unstable aminoacyl-AMPs, which are converted to aminoacyl-tRNAs or aminoacyl-peptidyl carrier proteins that can be used in protein synthesis or non-ribosomal peptide synthesis, respectively. In contrast, enzyme-generated adenylates have not been used as activated intermediates for *in vitro* protein bioconjugation or semisynthesis by bioorganic chemists^5,6^. Instead, the most widely adopted protein bioconjugation methods rely on reversible enzyme-catalyzed transpeptidation reactions^7–13^ or non-enzymatic pre-synthesis of activated precursors^14–16^ to provide a driving force for peptide bond formation.

The E1-like (ThiF) enzyme superfamily is comprised of structurally and mechanistically related proteins that share a common mechanistic step involving adenylation of a C-terminal α-carboxylate to generate a reactive peptidyl-*O*-AMP mixed anhydride electrophile^17,18^. Based on its capacity to react with diverse nucleophiles, this shared intermediate provides biological systems with access to protein and peptide C-terminal thioesters^3^, thiocarboxylates^19,20^, succinimides^21,22^ and (iso)peptide bonds^23,24^ that function in biological processes ranging from ubiquitination to biosynthesis of metabolites including thiamin, molybdopterin, and ribosomally synthesized and post-translationally modified peptide (RiPP) natural products. Despite their versatility, E1-like enzymes have not been developed for *in vitro* applications in peptide bond synthesis as many of the best studied E1-like enzymes require ubiquitin-like proteins as their substrates. In contrast to many other E1-like enzymes, *E. coli* MccB, which natively functions in formation of a C-terminal phosphoramidate linkage in biosynthesis of the RiPP natural product microcin C7, accepts a short heptapeptide as its substrate^21,25^. This feature makes MccB a promising candidate tool for protein bioconjugation. However, MccB’s native reaction chemistry does not couple C-terminal adenylation to a peptide bond formation reaction that would be useful for protein bioconjugation.

Here, we report the mechanism-guided design of an MccB-based toolbox for C-terminal activation and protein modification. We show that *E. coli* MccB has a latent capacity to catalyze C-terminal adenylation on non-native substrates. The resultant C-terminal peptidyl-*O*-AMP electrophile can react with a variety of exogenous nucleophiles, including hydrazines, alkoxyamines, amines, and thiols, to form diverse C-terminal functional groups. We find that MccB-generated thioesters can undergo *trans*-thiolation and *S*-to-*N* acyl transfer, analogous to the reactions involved in the eukaryotic ubiquitination cascade, that make them attractive intermediates for protein bioconjugation. We develop the <u>T</u>hio<u>e</u>sterification <u>C</u>-terminal <u>H</u>andle (TeCH-tag), a sequence that enables MccB-catalyzed, ATP-driven synthesis of protein C-terminal thioesters, an important class of protein semisynthesis intermediates that were previously only directly accessible via a method that relies on engineered self-splicing inteins. We apply the MccB/TeCH-tag system for high-yield, ATP-dependent protein bioconjugation via expressed protein ligation and enzyme-catalyzed expressed protein ligation. By mining the natural diversity of the MccB family, we develop two additional MccB/TeCH-tag pairs that are mutually orthogonal to each other and to the *E. coli* system, enabling the synthesis of more complex bioconjugates. Our method mimics the chemical logic of peptide bond synthesis that is widespread in biology for high-yield *in vitro* synthesis of protein bioconjugates that will advance our understanding of biological systems.

### Enzyme mechanism-guided design of a tool for C-terminal functionalization

*E. coli* MccB is a member of the E1-like enzyme superfamily that catalyzes conversion of the C-terminal Asn residue of its heptapeptide substrate, MccA (MRTGNAN), to an isoasparagine (isoAsn)-AMP phosphoramidate (**Fig. 1A, B**)^21,25^. As its first mechanistic step, MccB is proposed to catalyze C-terminal adenylation (or *O*-AMPylation) of MccA, producing an MccA-*O*-AMP intermediate that is captured by the β-carboxamido nitrogen group of the C-terminal Asn (N7) residue to form a succinimide intermediate (**Fig. 1C**). This succinimide is subsequently *N*-AMPylated and hydrolyzed to form the C-terminal isoAsn-AMP product^21^. This modified heptapeptide is further tailored by phosphoramidate aminopropylation^26,27^ and acts as a ‘Trojan horse’ antibiotic that is cleaved by an endogenous protease in target cells to form isoAsn-AMP, an aspartyl-adenylate mimic that inhibits the aspartyl-tRNA synthetase^28,29^. While most families within the E1-like superfamily act on ubiquitin-like β-grasp fold proteins with a C-terminal Gly residue^17,18^, MccB family enzymes are unique in that they accept as their substrates short, genetically encoded peptides terminating with C-terminal Asn (**Fig. 1A, B**)^30,31^ We hypothesized that if we substituted the C-terminal Asn of the MccA substrate with another amino acid, MccB would retain the ability to *O-*AMPylate the non-native substrate and that in the absence of a *cis* nucleophile in the substrate, the peptidyl-*O*-AMP electrophile could react with an exogenous nucleophile for C-terminal functionalization.

**Figure 1.**
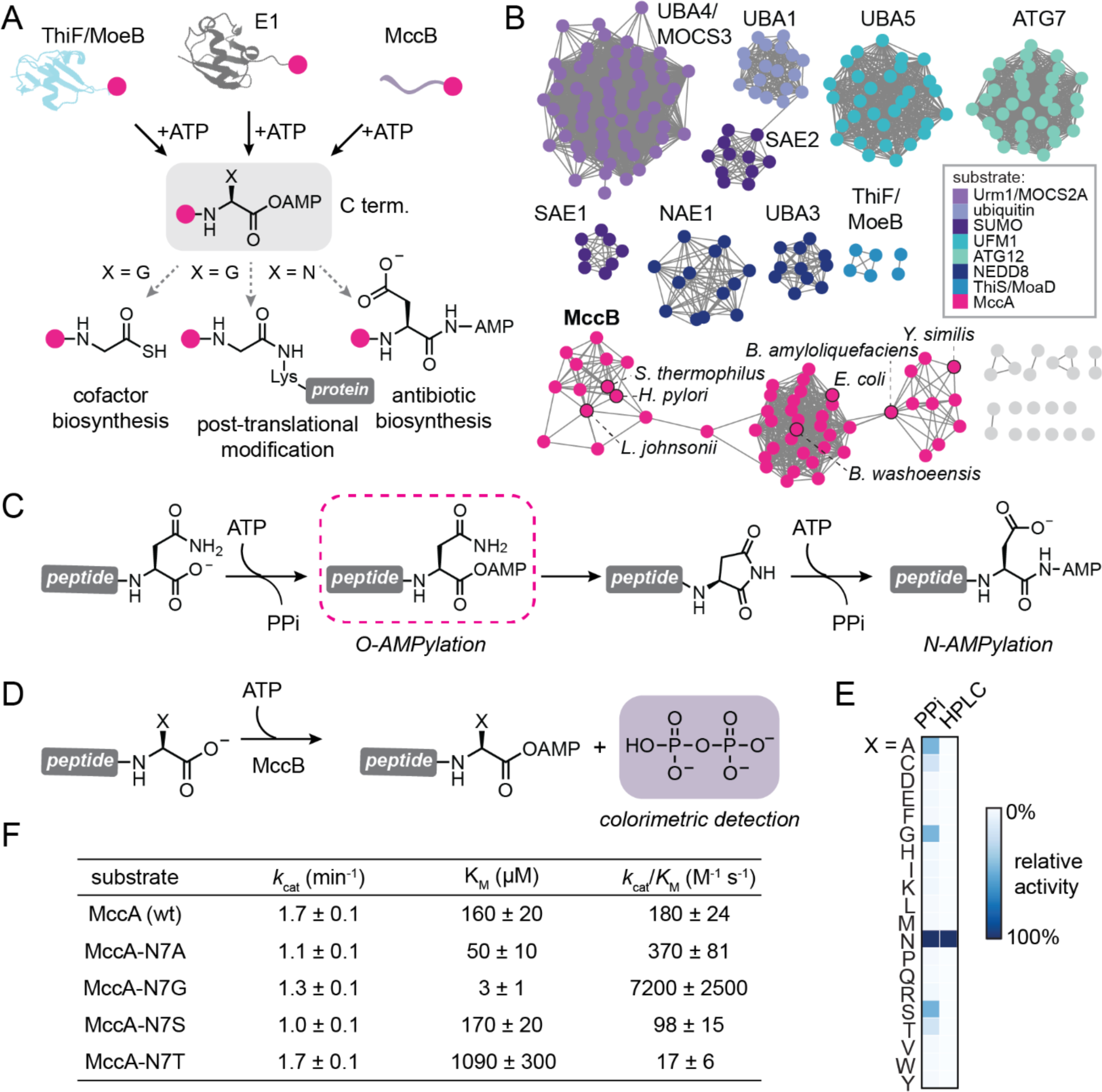
MccB catalyzes C-terminal *O*-AMPylation on non-native substrates. (A) MccB enzymes are members of the E1-like/ThiF superfamily, which have in common the formation of a C-terminal peptidyl-*O*-AMP intermediate. (B) Members of the E1-like superfamily that act on C-termini generally recognize ubiquitin-like proteins that terminate in Gly as their substrates; the MccB family is an exception as it acts on short peptides (MccAs) that terminate in Asn. (C) The reaction catalyzed by MccB in its native context. (D) The C-terminal residue of *E. coli* MccA was varied to the 19 non-native amino acids and MccB’s ability to catalyze C-terminal *O*-AMPylation was assayed using a colorimetric assay for pyrophosphate. (E) MccA-N7A, N7G, N7S, and N7T stimulate MccB-catalyzed pyrophosphate formation. (F) Steady-state kinetic parameters for MccB-catalyzed pyrophosphate release when wt MccA or MccA variants are used as substrates.

To test the hypothesis that MccB retains a latent capacity to *O*-AMPylate peptide substrates lacking the conserved C-terminal Asn residue, we synthesized a small library of non-native MccA peptides in which the C-terminal residue was varied to the 19 amino acids other than Asn (MccA-N7X). Because the peptidyl-*O*-AMP product of MccB-catalyzed *O*-AMPylation is expected to be hydrolytically unstable, we initially screened the ability of these MccA variants to stimulate ATP consumption by MccB using an enzyme-coupled assay to detect formation of the pyrophosphate (PP_i_) by-product (**Fig. 1D**)^32,33^. We observed substantial stimulation of PP_i_ formation when MccA-N7A, N7G, N7S, or N7T were used as substrates for MccB (**Fig. 1E**, **Fig. S1**). We determined the steady-state kinetic parameters for MccB-catalyzed pyrophosphate formation with each of these substrates and found that the catalytic efficiencies (*k*_cat_/*K*_M_) were similar to or exceeded the *k*_cat_/*K*_M_ measured for wild-type MccA (**Fig. 1F**, **Fig. S2**). We next tested whether this ATP consumption resulted in the formation of any stably AMPylated product using an HPLC-MS assay. While we observed a clear +329.0485 Da peak corresponding to AMPylation when wild-type MccA was used as a substrate, no stably AMPylated product was observed for the MccA-N7A, N7G, N7S, or N7T variants (**Fig. 2A, B**, **Fig. S3**). These results are consistent with a previous study that showed that N7 is required for microcin C7’s antimicrobial activity^34^. Together these data raise two possibilities: 1) Binding of MccA-N7X variants to MccB stimulates unproductive ATP hydrolysis; or 2) MccA-N7X variants bind to MccB and are *O*-AMPylated, but the resulting mixed anhydride is hydrolytically unstable and cannot be detected by HPLC-MS.

**Figure 2.**
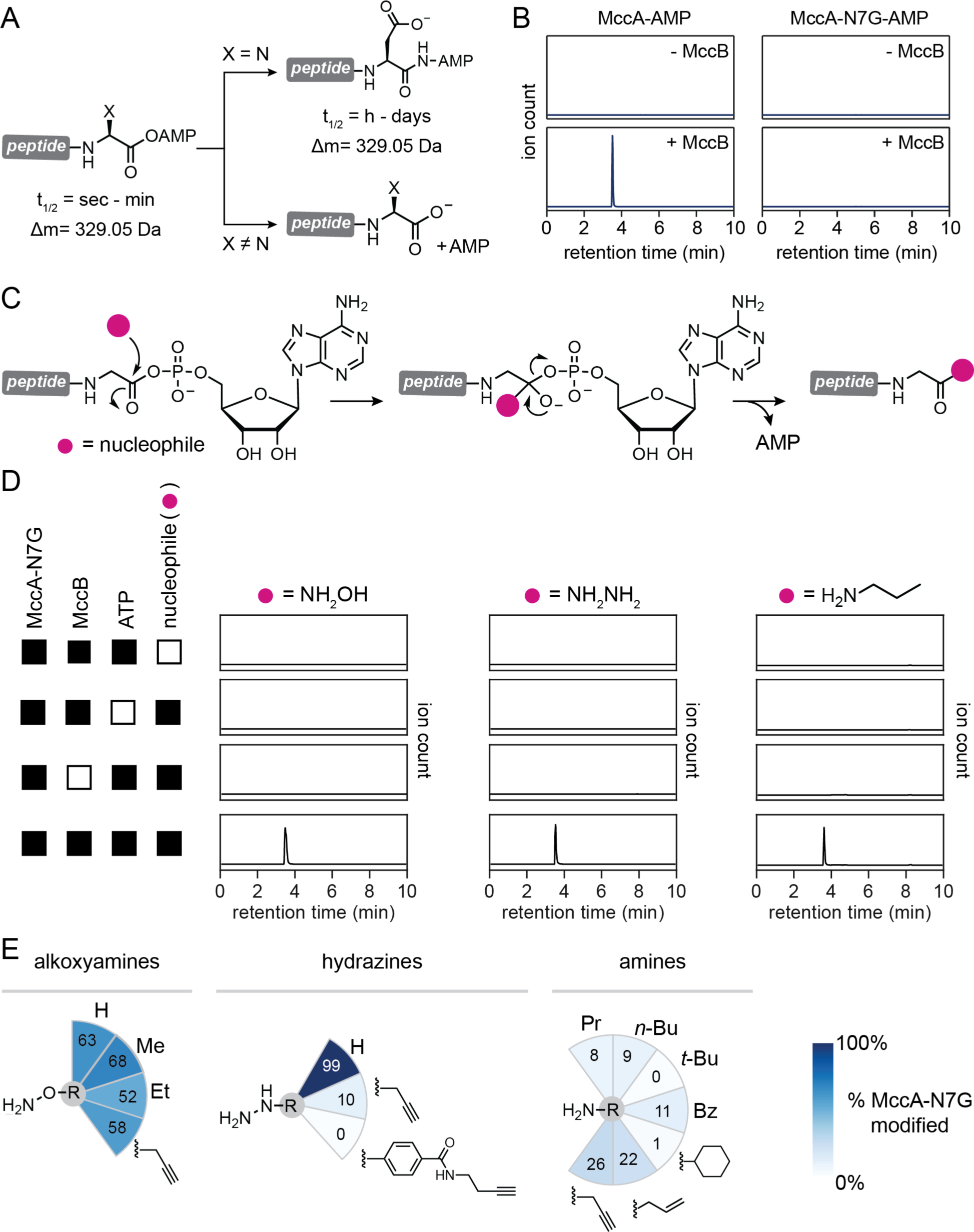
MccB catalyzes formation of a peptidyl-*O*-AMP intermediate that can react with exogenous nucleophiles. (A) MccA-*O*-AMP undergoes reaction with its C-terminal Asn side chain to form a succinimide intermediate and subsequently a stably *N*-AMPylated product that can be detect by LC-MS (top). In contrast, MccA-N7G-O-AMP is expected to undergo hydrolysis on a timescale incompatible with LC-MS detection. (B) LC-MS analysis of MccB reactions with wild-type MccA (left) or MccA-N7G (right) as a substrate. (C) Hypothesized MccA-N7G-*O*-AMP reactivity with exogenous nucleophiles. (D) LC-MS analysis of MccB-catalyzed reactions containing ATP (5 mM), MccA-N7G (250 μM), and hydroxylamine (150 mM, left), hydrazine (150 mM, middle), or propylamine (150 mM, right) reveals that MccA-N7G-*O*-AMP can react with exogenous nucleophiles. (E) Radial heatmaps show the percent conversion of MccA to nucleophile-modified MccA for a panel of alkoxyamine, hydrazine, and amine nucleophiles.

We hypothesized that if an electrophilically activated *O*-AMPylated C terminus is formed, an exogenously added nucleophile, hydroxylamine, could compete with water to attack the electrophilic group, resulting in MccA-N7X bearing a C-terminal hydroxamate that could be distinguished from the unmodified peptide based on its mass (+15.0109 Da) (**Fig. 2C**). Such experiments have been used previously to detect electrophilic enzymatic intermediates including anhydrides and thioesters^35,36^. To test this hypothesis, we included 150 mM hydroxylamine in reactions containing MccB, ATP, and MccA-N7G (m/z 706.3306). HPLC-MS analysis of these reactions revealed formation of a new peak corresponding to formation of a C-terminal hydroxamate (m/z 721.3415) that was dependent on MccB, ATP, and hydroxylamine (**Fig. 2D**). Similar results were obtained using hydrazine as the trapping nucleophile (**Fig. 2D**, Fig**. S4**). These data support the hypothesis that MccB catalyzes *O*-AMPylation of non-Asn C-terminal residues and provide direct evidence for the *O*-AMPylated intermediate in the proposed mechanism of MccB.

We next sought to define features of the nucleophile that impact the efficiency of capture of the *O*-AMPylated intermediate. We initially screened a panel of hydroxylamines, hydrazines, and amines by measuring their ability to modify the C terminus of MccA-N7G in an MccB-dependent manner using HPLC-MS (**Fig. 2E**, **Fig. S4**). Among hydroxylamine nucleophiles, increasing the size of the *O*-substituent decreased the efficiency of C-terminal modification. Similarly, we found that a recently reported phenylhydrazine-based nucleophilic probe (N-(but-3-yn-1-yl)-4-(2-hydrazineylethyl)benzamide)^37^ was unable to modify the C terminus of MccA-N7G. These results suggest that nucleophile size is an important factor in the efficiency of MccB-catalyzed MccA-N7G modification and raise the hypothesis that the intermediate is not released from the enzyme but reacts with exogenous nucleophiles within the enzyme active site. In contrast to the efficient capture observed with hydroxylamine and hydrazine, small primary and secondary amines modified MccA inefficiently (0-26%) (**Fig. 2DE**, **Fig. S4**). These results suggest that nucleophile strength is also an important factor in efficient capture of the *O*-AMPylated intermediate.

### Engineered MccB reactivity mimics the eukaryotic ubiquitination cascade for C-terminal modification

Inspired by the relationship between MccB and E1 ubiquitin-activating enzymes, we next sought to examine whether *O*-AMPylated MccA-N7G could react with thiol nucleophiles to form C-terminal thioesters. In the canonical ubiquitination cascade, an E1 catalyzes C-terminal *O*-AMPylation of ubiquitin (Ub), activating Ub for nucleophilic attack by a Cys side chain to generate an E1-bound Ub C-terminal thioester (denoted E1∼Ub). E1∼Ub then undergoes *trans*-thiolation with a Cys side chain of an E2 ubiquitin-conjugating enzyme to form E2∼Ub. Finally, an E3 ubiquitin ligase binds both E2∼Ub and the substrate to catalyze *S*-to-*N* acyl transfer to a Lys side chain from the substrate to form an isopeptide bond (**Fig. 3A**)^38^. We hypothesized that capture of MccB-generated MccA-N7G-*O*-AMP with a thiol nucleophile would enable both C-terminal thioester synthesis and subsequent reactions, including *trans*-thiolation and *S*-to-*N* acyl shift, that mimic the ubiquitination cascade.

**Figure 3.**
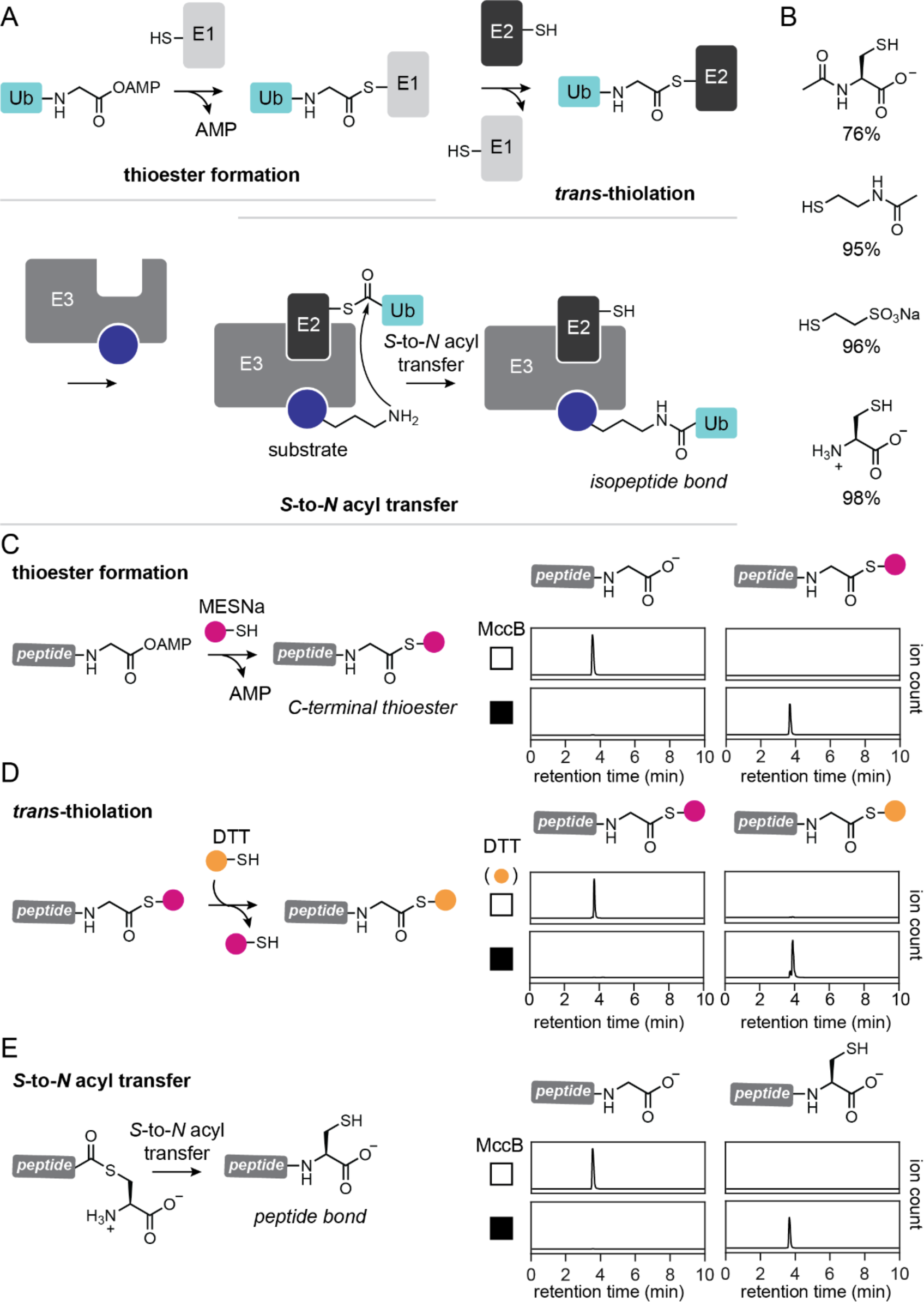
Engineered MccB reactivity mimics the eukaryotic ubiquitination cascade for C-terminal protein modification. (A) The ubiquitination cascade involves C-terminal thioesterification, *trans*-thiolation, and S-to-N acyl shift to transfer ubiquitin to substrate residues. (B) Efficient capture of MccA-N7G-*O*-AMP with thiol nucleophiles. (C-E) Engineered MccB reactivity initiated by (C) C-terminal thioesterification enabled (D) *trans*-thiolation and (E) *S*-to-*N* acyl transfer, mimicking the reactions of the ubiquitination cascade.

We first tested whether MccB could catalyze ATP-dependent C-terminal thioesterification of MccA-N7G analogous to E1-catalyzed formation of the E1∼Ub thioester. As a surrogate for a Cys side chain, we used *N*-acetyl-L-cysteine as the thiol nucleophile and found that the C terminus of MccA-N7G was modified in an MccB- and ATP-dependent manner in 76% yield within 16 hours (**Fig. 3B**, **Fig. S4**). We then tested a larger panel of thiol nucleophiles, including *N*-acetylcysteamine, DTT, sodium 2-mercaptoethane sulfonate (Mesna), and 4-mercaptophenylacetic acid (MPAA) (**Fig. 3C**, **Fig. S4**). With the exception of MPAA, these thiol nucleophiles modified MccA-N7G in >95% yield. We attribute the inability of MPAA to modify MccA-N7G to the constraints on nucleophile size and structure that are imposed by the enzyme active site. These results demonstrate that the MccA-N7G-*O*-AMP intermediate can be efficiently transferred to thiol nucleophiles, mimicking the E1-catalyzed step in the ubiquitination cascade.

The next step in the ubiquitination cascade involves *trans*-thiolation between E1∼Ub and a Cys side chain on an E2 enzyme^39^. To test whether MccB-generated thioesters could undergo *trans*-thiolation, we initially used MccB to generate an MccA-N7G Mesna thioester. We then tested whether this thioester could exchange with a second thiol, DTT (**Fig. 3D**, **Fig. S5**). First, we used MccB to generate the MccA-N7G Mesna thioester in a 16 h reaction with 5 mM Mesna. We then added DTT to 15 mM to establish a molar excess. We observed complete conversion of the Mesna thioester to the DTT thioester over 4 hours, demonstrating that MccB thioesters can undergo *trans*-thiolation (**Fig. 3D**). To examine whether the exchange reaction is enzyme-catalyzed or occurs non-enzymatically in solution, we next tested the ability of the Mesna thioester to undergo *trans*-thiolation with MPAA, a thiol nucleophile that is unable to participate in the initial enzyme-catalyzed thioesterification reaction. We found that MPAA functioned efficiently in *trans*-thiolation (**Fig. S6**), suggesting that this reaction occurs non-enzymatically in solution.

The final step in ubiquitin transfer to a substrate protein involves an E3 ubiquitin ligase-catalyzed *S*-to-*N* acyl shift in which Ub is transferred from the E2 Cys residue to a substrate Lys side chain^40^. To examine whether MccB-generated thioesters can undergo *S*-to-*N* acyl shift, we used Cys as a nucleophile for capture of MccA-N7G-*O*-AMP. We observed complete modification of MccA-N7G by Cys as indicated by the disappearance of the MccA-N7G peak and appearance of a +103.0083 Da peak corresponding to addition of a Cys residue (**Fig. 3E**, **Fig. S4**). To distinguish whether this peak resulted from formation of a thioester with the Cys β-thiol or formation of an amide bond with the α-amine, we tested whether MccA-N7G-Cys could be modified by the thiol-reactive compound maleimide. We observed complete conversion to a maleimide-modified species (+200.0034 Da, **Fig. S7**), indicating that *S*-to-*N* acyl shift and amide bond formation had occurred.

In addition to their function in the installation of Ub and related protein modifications onto protein substrates in biological systems, *S*-to-*N* acyl transfer reactions have been widely employed for convergent synthesis of chemically tailored proteins via native chemical ligation (NCL)^41,42^. In NCL, a peptide modified with a C-terminal thioester can condense with a peptide bearing an N-terminal Cys residue to form a native peptide bond through a mechanism involving *trans*-thiolation between the Cys side chain and thioester followed by *S*-to-*N* acyl shift^42^. While the initial *trans*-thiolation step in NCL is reversible, the *S*-to-*N* acyl shift that forms the amide bond is irreversible, driving the reaction to high yield. Inspired by the utility of this reaction, we sought to test whether N-terminal Cys peptides could similarly capture MccA-N7G-*O*-AMP and undergo *S*-to-*N* acyl shift to enable a peptide ligation reaction. We first tested a Cys-Trp dipeptide nucleophile and observed no modification of MccA-N7G, suggesting that the larger nucleophile is unable to access MccA-N7G-*O*-AMP within the MccB active site (**Fig. S8**). However, when we included both Mesna and Cys-Trp in the reaction, MccA-N7G was converted to MccA-N7G-CW in 82% yield in 4 hours (**Fig. S8**). This result suggests that MccA-N7G can undergo MccB-catalyzed thioesterification with Mesna, followed by *trans*-thiolation and *S*-to-*N* acyl shift with Cys-Trp in one pot, enabling enzyme-catalyzed NCL starting from an unactivated peptide with a C-terminal carboxylate.

### Applying MccB for protein C-terminal thioesterfication and peptide bond formation

Encouraged by the ability of MccB to catalyze MccA-N7G thioesterification to activate it for C-terminal peptide ligation, we hypothesized that MccA-N7G could be developed as a tag for C-terminal protein activation. Previous functional characterization of MccB demonstrated that an N-terminal maltose binding protein (MBP) fusion of MccA is recognized as a substrate by MccB and can be modified with an N-P bond to AMP^31^. To test whether MccA-N7G could similarly enable protein C-terminal activation and peptide ligation, we fused GFP to MccA-N7G (a sequence that we term the <u>T</u>hio<u>e</u>sterification <u>C</u>-terminal <u>H</u>andle, or TeCH-tag) (**Fig. 4A**, **Fig. S9**). We first tested the ability of MccB to catalyze ATP-dependent thioesterification of GFP-TeCH-tag with 1 mM or 5 mM Mesna. In the presence of 5 mM Mesna, we observed near-quantitative conversion of GFP-TeCH-tag to the Mesna thioester with 30 min, while 1 mM Mesna gave >75% conversion with somewhat slower kinetics (**Fig. 4B**, **Fig. S9**). The MccB/TeCH-tag system therefore provides and efficient method to generate C-terminal protein thioesters.

**Figure 4.**
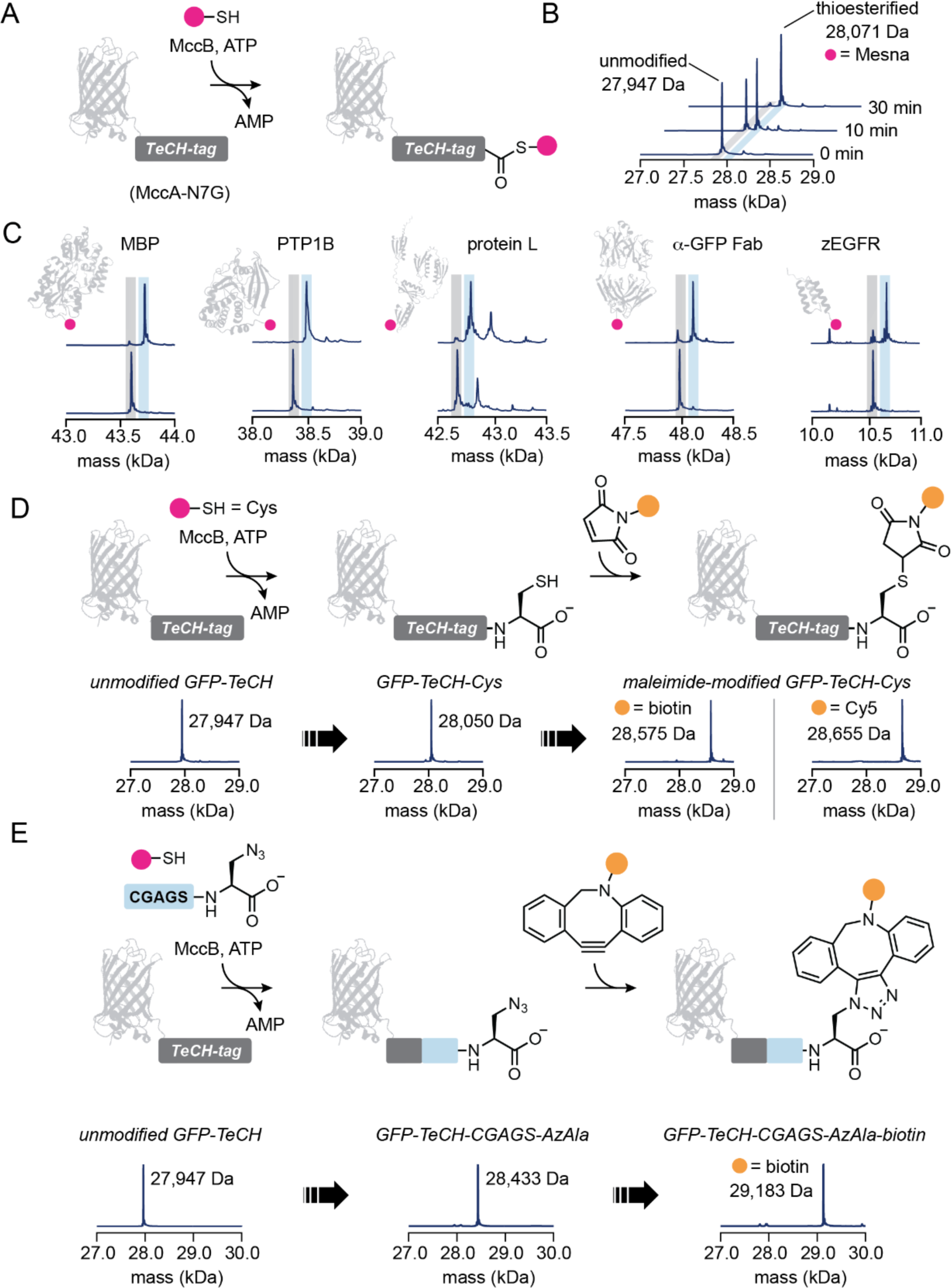
Fusion of the <u>T</u>hio<u>e</u>sterification <u>C</u>-terminal <u>H</u>andle (TeCH-tag) to proteins enables MccB-catalyzed, ATP-dependent formation of C-terminal thioesters. (A) Fusion of the TeCH-tag to GFP for C-terminal thioesterification. (B) MccB catalyzes ATP-dependent thioesterification of GFP-TeCH-tag within 30 min. (C) MccB catalyzes C-terminal thioesterification of TeCH-tag fusions of MBP, the catalytic domain of protein tyrosine phosphatase 1B (PTP1B_1-321_), protein L, an α-GFP recombinant antibody, and an EGFR-targeting affibody. The * indicates an α-gluconylated form of protein that is an artifact of His-tag purification. (D) MccB catalyzes C-terminal thioesterification of GFP-TeCH-tag with cysteine, with subsequence *S*-to-*N* acyl shift leading to formation of a peptide bond as evidence by the maleimide reactivity of the bioconjugate. (E) GFP-TeCH-tag can be modified by expressed protein ligation via a Mesna thioester intermediate reaction in a one-pot reaction with MccB, ATP, Mesna, and the N-terminal Cys peptide CGAGS-azidoalanine.

To examine the general utility of the MccB/TeCH-tag system, we appended the TeCH-tag to a diverse panel of proteins, including MBP; the catalytic domain (residues 1-321) of protein-tyrosine phosphatase 1B (PTP1B_1-321_); an anti-GFP recombinant antibody (α-GFP rAb); protein L; and an endothelial growth factor receptor (EGFR)-targeting affibody (zEGFR). We treated these TeCH-tag fusion proteins with MccB (5 μM), ATP (5 mM), and Mesna (5 mM) and found that they were thioesterified in near-quantitative yield within 1-16 hours (**Fig. 4C**). While GFP, MBP, and PTP1B_1-321_ were completely converted to thioester in 1 h, zEGFR required 6 h, and protein L and α-GFP rAb required 16 h. Notably, the α-GFP rAb, based on a scaffold derived from the therapeutic antibody Trastuzumab^43^, contains five disulfide bonds linking its light and heavy chains. We found that the rAb could be thioesterified in quantitative yield under conditions that omit reducing agent, keeping the disulfide bonds required for antibody function intact. These results demonstrate that the MccB/TeCH-tag system is broadly applicable for generating protein C-terminal thioesters.

C-terminal protein thioesters are key intermediates in expressed protein ligation (EPL), a powerful protein semisynthesis method that involves NCL between a recombinantly expressed protein C-terminal thioester and a synthetic N-terminal Cys peptide to form a native peptide bond^44^. Prior applications of EPL have relied on fusion of the protein to be modified with an engineered intein to generate the protein C-terminal thioester^6^. We examined whether the MccB/TeCH-tag system could be deployed to expand the toolbox for C-terminal thioester formation in the context of EPL. We initially examined whether MccB could catalyze incorporation of single Cys residue at the C terminus of a Cys-free variant of GFP (cfGFP)^45^ via an amide bond. We incubated cfGFP with MccB (5 μM), ATP (5 mM), and Cys (5 mM) and found that Cys was ligated to cfGFP in in 99% yield in 1 h (**Fig. 4D**, **Fig. S10**). Following Cys incorporation, the Cys side chain was quantitatively modified with biotin-maleimide or Cy5-maleimide, demonstrating that *S*-to-*N* acyl shift had occurred to produce a free Cys side chain.

We next tested whether a workflow involving thioesterification with Mesna, *trans*-thiolation with an N-terminal Cys peptide, and *S*-to-*N* acyl shift could be applied for ligation of cfGFP to a synthetic N-terminal Cys peptide bearing an azide functional group. We initially tested this scheme by incubating cfGFP with MccB (5 μM), ATP (5 mM), Mesna (5 mM), and a Cys-Trp dipeptide (5 mM). We observed complete conversion of the cfGFP-Mesna thioester to cfGFP-CW within 3 h (**Fig. S11**). We next performed a reaction in which TeCH-tagged cfGFP was incubated with MccB, ATP, Mesna, and peptide with the sequence CGAGS-3-azido-L-Ala. We found that cfGFP-TeCH-tag could be quantitatively modified with the azide-bearing peptide (**Fig. 4E**). The azide modified protein could then undergo strain-promoted azide-alkyne cycloaddition (SPAAC) with dibenzocyclooctyne (DBCO)-biotin (**Fig. 4E**). MccB can thus serve as a catalyst for protein C-terminal thioesterification to enable EPL.

Although EPL has been a powerful and widely adopted tool for protein semisynthesis, it has been limited by the need for Cys at the ligation junction, requiring the introduction of a nucleophilic, redox-active amino acid that can affect downstream applications. Several methods have been developed for removal of the thiol group following ligation, but they are generally only applicable to proteins that lack other Cys residues^46,47^. To overcome this limitation, a related method termed enzyme-catalyzed EPL has been developed to increase flexibility in terms of the sequence at the ligation junction^48^. In enzyme-catalyzed EPL, a C-terminal protein thioester is used as a substrate for the engineered peptide ligase subtiligase^14,49^, which has broad N-terminal specificity and eliminates the requirement for Cys at the ligation site. While this method has been applied to study phosphoregulation of the tyrosine phosphatase PTEN by introduction of phosphoresidues at the specific sites^48,50^, yields were limited by a competing subtiligase-catalyzed thioester hydrolysis reaction. We hypothesized that application of MccB for ATP-dependent thioester generation in this context would drive yields higher because it would enable thioester regeneration from the inactivated hydrolysis product (**Fig. 5A**).

**Figure 5.**
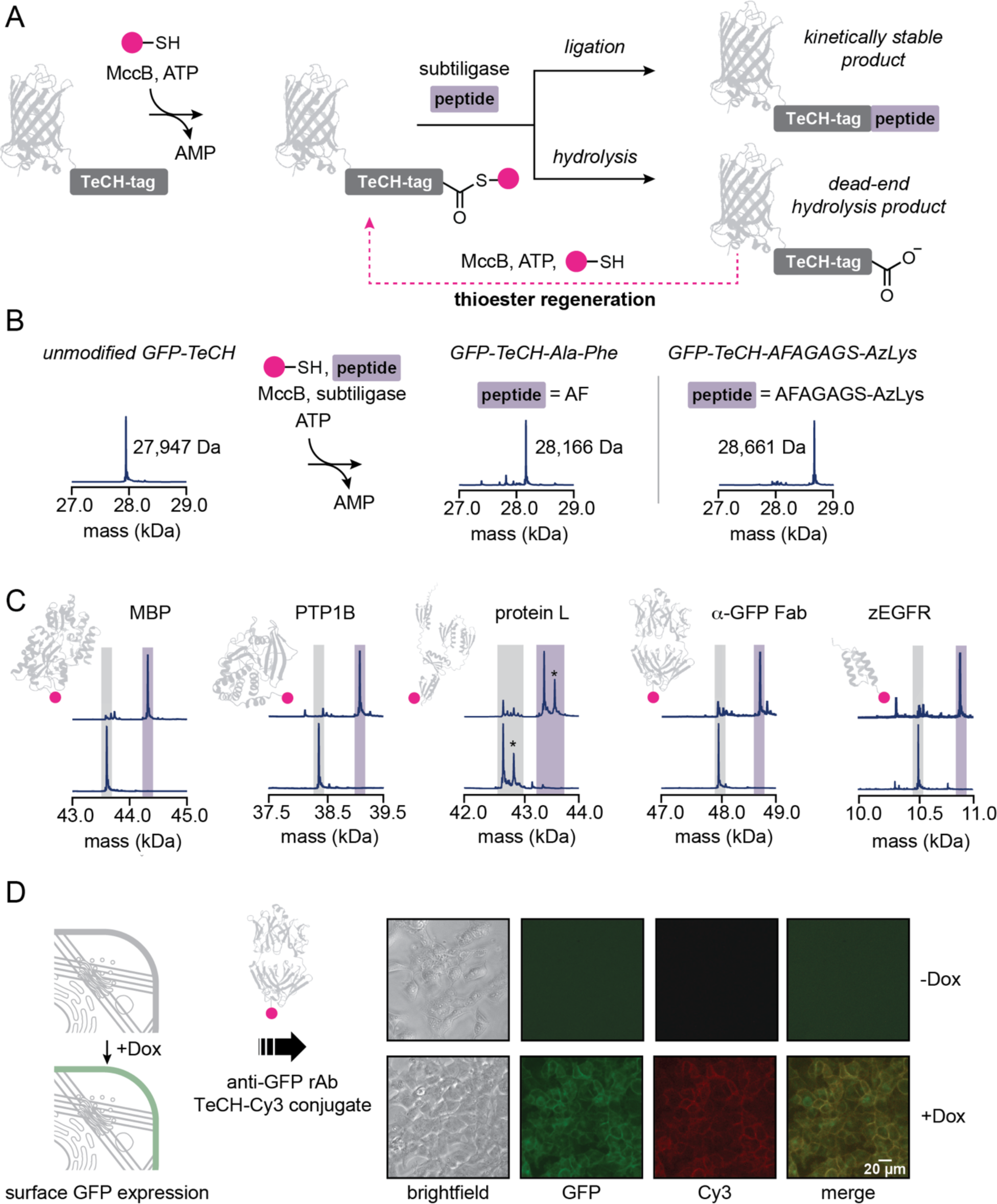
MccB enables ATP-dependent thioester formation and regeneration for enzyme-catalyzed expressed protein ligation. (A) Enzyme-catalyzed expressed protein ligation is limited by subtiligase-catalyzed hydrolysis of the thioester substrate, generating a dead-end product. We used MccB for ATP-dependent thioester formation and regeneration, enabling reactivation of the dead-end hydrolytic product. (B) High yield one-pot MccB- and subtiligase-catalyzed ATP-dependent peptide ligation to GFP-TeCH. (C) MccB and subtiligase catalyze ATP-dependent peptide ligation to TeCH-tag fusions of MBP, the catalytic domain of protein tyrosine phosphatase 1B (PTP1B_1-321_), protein L, an α-GFP recombinant antibody, and an EGFR-targeting affibody. The * indicates an α-gluconylated form of protein that is an artifact of His-tag purification. (D) MccB- and subtiligase-catalyzed peptide ligation and strain promoted azide-alkyne cycloaddition were used to synthesizea-GFP rAb-Cy3 for staining of a HEK293T cell line engineered for doxycycline-inducible expression of cell surface GFP.

To test this hypothesis, we first incubated TeCH-tagged GFP with MccB (5 μM), ATP (5 mM), Mesna (5 mM), subtiligase (5 μM), and the dipeptide Ala-Phe (5 mM) (**Fig. 5B**, **Fig. S12**). We found that GFP was ligated to Ala-Phe in near-quantitative yields within 4 h. We next tested MccB/subtiligase-catalyzed ligation of GFP to a longer peptide, AFAGAGS-azidolysine (5 mM), which contains an azide for downstream modification using click chemistry. We found that GFP could also be modified efficiently with this peptide (**Fig. 5B**). To test whether the efficiency of the ligation reaction depends on the protein substrate, we tested the reaction on our panel of TeCH-tag fusion proteins (MBP, PTP1B_1-321_, α-GFP rAb, protein L, and zEGFR) (**Fig. 5C**). We found that all of the proteins tested could be efficiently ligated to AFAGAGS-azidolysine in high yields, highlighting the general utility of MccB for driving subtiligase-catalyzed peptide ligation.

In contrast to the high yields obtained using the MccB/TeCH-tag system, previous enzyme-catalyzed EPL experiments achieved only ∼30-50% yield using ∼9 mM (10 mg/mL) of the peptide to be ligated in the reaction^48^. Investigation of the Ala-Phe concentration dependence of the MccB/subtiligase-catalyzed ligation reaction showed that high yields (83%) were maintained even when the Ala-Phe concentration was reduced to 1 mM, with ∼50% yield maintained at 0.5 mM Ala-Phe (**Fig. S12**). Unexpectedly, we observed that when Ala-Phe was omitted from the reaction, a substantial fraction of GFP was converted to a species with a mass of −18±1 Da compared to unmodified GFP. Appearance of this species is dependent on the presence of both MccB and subtiligase. We hypothesize that this product results from MccB- and subtiligase-catalyzed cyclization of GFP in which the C-terminal thioester is ligated to the N terminus of the same protein molecule. Although MccB- and subtiligase-catalyzed protein cyclization could be a useful addition to protein chemistry toolbox, we hypothesize that undesired cyclization could be avoided by careful selection of the N-terminal residue of the protein of interest and the subtiligase variant employed for thioester ligation such that N terminus is a poor substrate for the chosen subtiligase variant^51^.

To test the utility of bioconjugates synthesized with MccB/TeCH-tag enzyme-catalyzed EPL method in a biological context, we constructed a HEK293T cell line that expresses cell surface GFP under the control of a tetracycline/doxycycline-inducible promoter. We synthesized a Cy3-modified α-GFP rAb by using MccB and subtiligase to ligate AFAGAGS-azidolysine onto the C-terminal TeCH-tag fused to the heavy chain, followed by SPAAC with DBCO-Cy3 (**Fig. S13**). We then stained doxycycline (Dox)-induced cells and uninduced cells with the α-GFP rAb-Cy3 conjugate. We observed robust Cy3 staining that colocalized with GFP in the Dox-induced cells, while both GFP signal and Cy3 signal were not observed in uninduced cells (**Fig. 5D**). These results demonstrate the utility of MccB- and subtiligase-catalyzed bioconjugation for incorporating probes into antibodies while maintaining their ability to bind their targets.

### Natural sequence diversity encompasses orthogonal MccA/MccB pairs

Epitope-specific bioconjugation enzymes enable modification of proteins to probe their functions, to discover inhibitors and drugs, to immobilize them for catalysis, and conjugate them to cytotoxic drugs, among many other applications^8,10,52^. To examine the sequence specificity of MccB for the MccA substrate, we synthesized a library of peptides in which each of the amino acid in each position of MccA was varied to the 19 non-native canonical amino acids (**Fig. 6A**, **Fig. S14**). We then measured formation of the C-terminally *N*-AMPylated phosphoramidate product for each of the MccA variants to query the stringency of MccB’s sequence specificity. We found that only wild-type MccA was fully converted to product over 16 h. Consistent with our previous results, no product formation was observed if the seventh position was varied to an amino acid other than Asn. Outside the C-terminal residue, substitutions in the first two positions of MccA had the largest effect on MccB activity, with significant product formation observed only for the M1W, M1Y, R2K, and R2L variants. At positions 3-6, significant product formation was observed for 6-8 different substitutions in each position. However, none of these variants were converted to product in quantitative yield despite the long reaction time, suggesting that MccB is an epitope-specific enzyme. Our data are broadly consistent with a previous study that examined the effect of substituting MccA positions 2-7 on microcin C7 production and antimicrobial activity *in vivo* and found that positions 4-6 are most tolerant of substitutions^31^.

**Figure 6.**
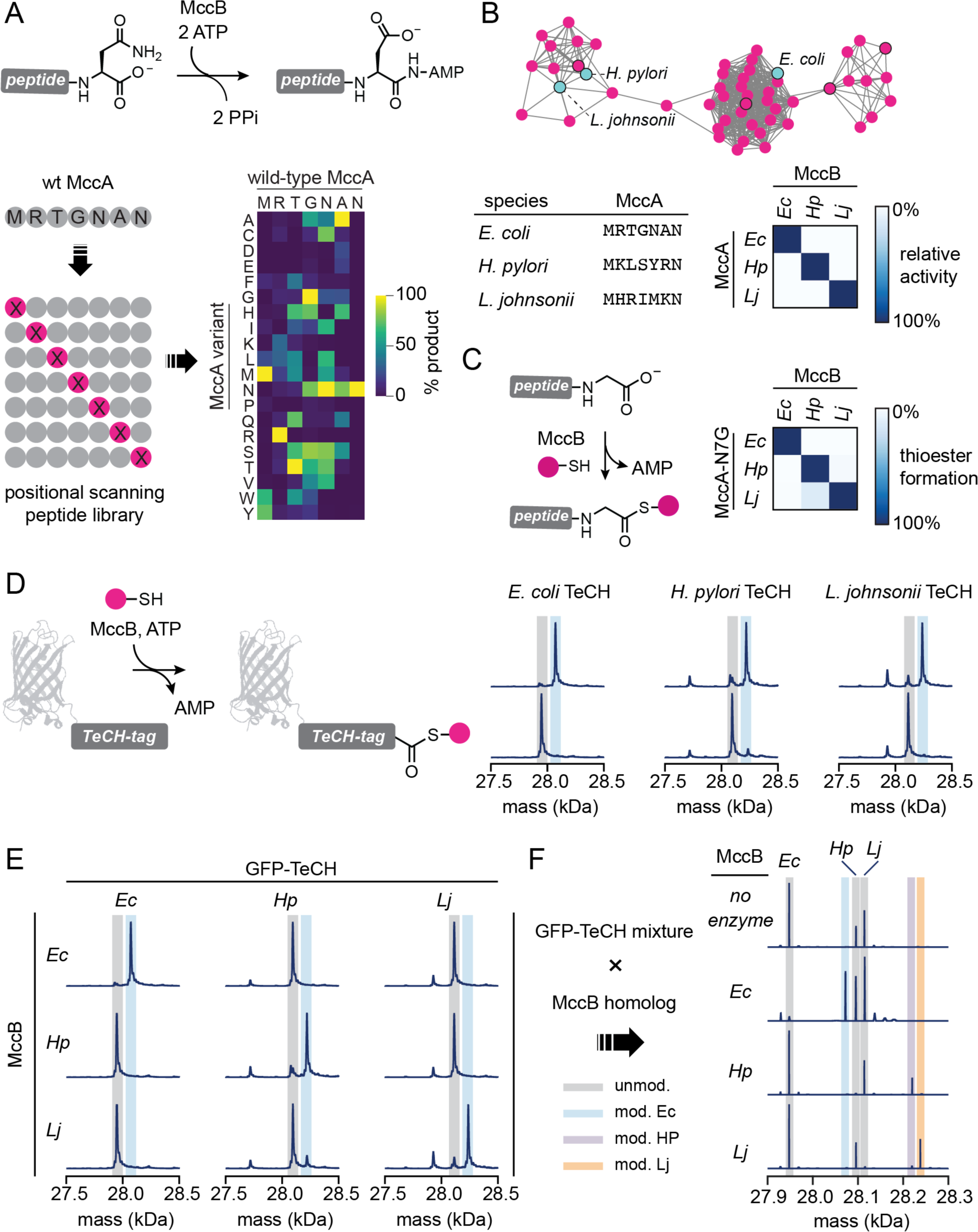
Natural MccA/MccB diversity encompasses orthogonal enzymes for C-terminal protein modification. (A) A positional scanning peptide library revealed that MccB is an epitope-specific enzyme. (B) The MccB enzyme family harbors numerous homologs that act on substrates distinct from *E. coli* MccA. Enzymes from *H. pylori* and *L. johnsonii* are highlighted in cyan. The *E. coli*, *H. pylori*, and *L. johnsonii* MccB were found to be mutually orthogonal enzymes. (C) The *E. coli*, *H. pylori*, and *L. johnsonii* enzymes catalyze C-terminal thioesterification of their respective MccA-N7G sequences and are orthogonal to one another. (D) LC-MS analysis of GFP-EcTeCH, GFP-HpTeCH, and GFP-LjTeCH shows that MccA-N7Gs from *E. coli*, *H. pylori*, and *L. johnsonii* can be deployed as TeCH-tags for C-terminal protein modification. (E) MccBs from *E. coli*, *H. pylori*, and *L. johnsonii* modify only proteins tagged with their cognate TeCH-tags. (F) In a mixture of GFPs fused to TeCH-tags from *E. coli*, *H. pylori*, and *L. johnsonii*, the MccB homologs from *E. coli*, *H. pylori*, and *L. johnsonii* are able to selectively modify their cognate TeCH-tags.

The application of epitope-specific bioconjugation enzymes can be limited by the relatively small number of available modification epitopes that restricts their use in the synthesis of more complex bioconjugates and in orthogonally targeting multiple different proteins in a mixture^10,52,53^. As a result, there is strong interest in identifying or engineering orthogonal enzyme/substrate pairs for protein modification. We hypothesized that the natural diversity of MccA and MccB might encompass mutually orthogonal enzyme-substrate pairs. Bioinformatic analyses have revealed that many bacterial genomes that encode MccB homologs also encode MccA-like peptides in the same gene cluster^30,31^. These analyses annotated 31 distinct, previously unknown heptapeptide MccA sequences as well as 14 longer putative MccAs. Based on the stringent sequence specificity of *E. coli* MccB, we hypothesized that the MccB homologs that recognize these distinct substrate sequences might be orthogonal to *E. coli* MccB and to one another. To test this hypothesis, we first measured the activity of the MccBs from *Helicobacter pylori* (HpMccB) and *Lactobacillus johnsonii* (LjMccB) toward their native substrates using an enzyme-coupled assay to measure the rate of PP_i_ release (**Fig. 6B**, **Fig. S15**). We found that both enzymes catalyzed PP_i_ release in the presence of their native substrates. We next tested whether *E. coli* MccB (EcMccB), HpMccB, and LjMccB had activity on the MccA substrates from the other species. We found that each enzyme only catalyzed efficient PP_i_ release in the presence of its cognate substrate, with <1% activity non-cognate sequences compared to cognate sequences (**Fig. 6B**, **Fig. S15**). This suggested that EcMccB/EcMccA, HpMccB/HpMccA, and LjMccB/LjMccA might be useful as mutually orthogonal enzyme-substrates pairs.

We next sought to test whether HpMccB and LjMccB could catalyze C-terminal thioesterification of HpMccA-N7G and LjMccA-N7G, respectively. We incubated each enzyme with its cognate MccA-N7G, ATP, and Mesna. Using LC-MS analysis, we found that each enzyme converted its cognate substrate to the C-terminal thioester in >97% yield within 16 h (**Fig. 6C**, **Fig. S16**, **Fig. S17**). We next tested whether EcMccB, HpMccB, and LjMccB catalyzed C-terminal thioesterification of MccA-N7Gs from other species. We did not detect thioesterification of non-cognate substrates by the EcMccB or LjMccB over 16 h, while a small of amount of thioesterification (13%) of LjMccA catalyzed by HpMccB was observed (**Fig. 6C**, **Fig. S17**). These results suggest that these three enzyme-substrate pairs possess a high degree of mutual orthogonality in terms of their ability to catalyze C-terminal thioesterification.

We also performed control experiments to measure the ability of each enzyme to catalyze formation of the *N*-AMPylated product from cognate and non-cognate MccAs (**Fig. S18**). Interestingly, and in contrast to our findings with MccA-N7G peptides, we observed substantial accumulation of non-cognate *N*-AMPylated products at the 16 h but not the 1 h timepoint. While the *L. johnsonii* and *H. pylori* systems were both mutually orthogonal with *E. coli* system, HpMccB catalyzed formation of LcMccA-*N*-AMP and LjMccB catalyzed formation of HpMccA-*N*-AMP. We hypothesize that this difference in orthogonality between the MccA and MccA-N7G substrates results from the higher efficiency of MccA-*O*-AMP capture in cis by the internal Asn carboxamide nucleophile compared to the exogenously added Mesna nucleophile. Supporting this hypothesis, we have observed that exogenous nucleophiles are unable to effectively compete with the internal β-carboxamido nucleophile when they are added to reactions with E. coli MccB and MccA. Our results are also consistent with previous studies that support a kinetic mechanism in which *O*-AMPylation is the rate-limiting step of the MccB-catalyzed reaction^21^.

Encouraged by our observation that EcMccB, HpMccB, and LjMccB are mutually orthogonal catalysts for MccA-N7G thioesterification, we fused HpMccA-N7G and LjMccA-N7G to GFP to test whether these sequences could be used for C-terminal protein thioesterification. We found that both enzymes converted their cognate TeCH-tagged GFPs to C-terminal thioesters, although in somewhat lower yield than was observed for the *E. coli* system. We hypothesized that incomplete conversion might be related to less efficient capture with Mesna and that this issue could be overcome by increasing the Mesna concentration, which we had previously set at 5 mM based on the requirements of the *E. coli* system. We found that increasing the Mesna concentration increased the thioester yield, with 50 mM Mesna supporting high-yield thioester formation (**Fig. 6D**).

To test the orthogonality of the three MccB/TeCH-tag systems that we developed, we examined the ability of each MccB to modify GFP-EcTeCH, GFP-HpTeCH, and GFP-LjTeCH. We found that each MccB was only able to modify its cognate GFP-TeCH (**Fig. 6E**). Next, we tested whether each MccB could selectively modify its cognate GFP-TeCH in a mixture of GFP-EcTeCH, GFP-HpTeCH, and GFP-LjTeCH (**Fig. 6F**). We observed that each of the MccBs only modified its cognate substrate, even at high (50 mM) Mesna concentrations. EcMccB/EcTeCH, HpMccB/HpTeCH, and LjMccB/LjTeCH therefore represent three new orthogonal C-terminal modification enzymes that greatly expand the available toolbox of epitope specific bioconjugation enzymes.

## Discussion

We designed the MccB/TeCH-tag system to mimic the chemical logic of peptide bond synthesis in biological systems to drive protein and peptide bioconjugation reactions to high yield. MccB/TeCH-tag can be used in the context of enzyme-catalyzed EPL for ATP-dependent thioester regeneration, driving the reaction equilibrium toward the desired ligation product and away from the dead-end thioester hydrolysis product formed by adventitious subtiligase reactivity. Our system avoids the reversibility of transpeptidases, such as sortase and asparaginyl endopeptidases, in which the desired ligation products are also transpeptidase substrates. Because this limitation is based on the position of the equilibrium, it cannot be overcome through transpeptidase engineering. Although depsipeptide (ester) and thiodepsipeptide (thioester) substrates have been used to drive transpeptidation, these substrates must be chemically synthesized, cannot be regenerated, and are mainly useful for N-terminal rather than C-terminal labeling.

The MccB/TeCH-tag system couples ATP cleavage to C-terminal activation via formation of a peptidyl-*O*-AMP for high-yield *in vitro* protein modification. Although acyl-*O*-AMPs are reactive electrophiles analogous to the acid chlorides and acid anhydrides often used in organic synthesis, they have not typically been viewed as modular intermediates that can be deployed for synthesis of modified proteins. We took advantage of our understanding of the enzymatic reaction mechanism of MccB to design a system for peptidyl-*O*-AMP synthesis that can be integrated with protein chemistry toolbox in modular fashion for bioconjugate synthesis. Introduction of the N7G substitution to MccA abolishes its ability to serve as a precursor to the antimicrobial compound microcin C7, but still supports the formation of a C-terminally *O*-AMPylated electrophile. We showed that MccA-N7G-*O*-AMP can react with alkoxyamines, hydrazines, amines, and thiols to form oximes, hydrazides, amides, and thioesters, respectively. We anticipate that this reactivity can easily be extended to other nucleophiles for synthesis of C-terminally modified proteins. For example, ammonia could be used as a nucleophile for modification of proteins by C-terminal amidation, which has recently been shown to target proteins for ubiquitin modification by SCF/FBXO31 and proteasomal degradation^54^. Other classes of nucleophiles such as alcohols could also capture the MccA-N7G-*O*-AMP intermediate for installation of protein C-terminal esters, which are present in prenylated proteins including Ras GTPases^55^.

We show that capture of MccA-N7G-*O*-AMP with thiol nucleophiles is particularly useful in protein bioconjugation as it converts the hydrolytically unstable peptidyl-*O*-AMP to a kinetically stable yet thermodynamically activated thioester. In biology, C-terminal thioesters function in enzymatic catalysis^56,57^, serve as intermediates in protein splicing^58,59^, and enable the installation of post-translational modifications including ubiquitin and ubiquitin-like proteins^3,18,40^. Although nature has evolved several strategies to generate C-terminal thioesters, only one, intein-mediated protein splicing, had previously been harnessed as a tool for producing recombinant protein C-terminal thioesters that serve as versatile intermediates for synthesis of chemically tailored proteins^6^. Recombinant C-terminal thioesters can be deployed for native chemical ligation to N-terminal Cys peptides in expressed protein ligation (EPL)^44^ or can be used as substrates for the engineered peptide ligase subtiligase in enzyme-catalyzed EPL^48^. These strategies have enabled precise manipulation of protein structure to advance our understanding of a broad range of biological questions^6^. The MccB/TeCH-tag system expands the toolbox for directly C-terminal thioester synthesis from unactivated protein α-carboxylates and therefore represents a broadly applicable tool for protein semisynthesis.

Our results indicate that MccB is an epitope-specific enzyme. Epitope-specific bioconjugation enzymes are valuable tools for protein modification, enabling installation of probes, payloads, and modifications that cannot be genetically encoded. In previous work, bacterial sortases have been widely applied for epitope-specific bioconjugation based on their selectivity for an LPXTG motif^12^. However, synthesis of complex bioconjugates can be limited by the relatively small number of available epitope-enzyme pairs, and few orthogonal sortase/sorting motif pairs have been identified in nature^60^. As a result, development of new orthogonal sortases has required intensive protein engineering efforts^52,53^. In contrast, the natural diversity of MccA/MccB pairs enabled us to readily generate two orthogonal tools for epitope-specific protein bioconjugation. Although we characterized three MccB/TeCH-tags, there are at least 31 distinct annotated heptapeptide MccA sequences with potential utility for protein bioconjugation^30,31^. Future genome mining or experimental approaches to identify orphan MccB substrates have the potential to further expand the number of available MccB/TeCH-tag systems and may enable the design of tailor-made bioconjugation enzymes that recognize user-defined sequences.

Advances in our ability to construct modified proteins have expanded the frontiers of our understanding of how post-translational modifications regulate transcription^61^, transduce cellular signals^62,63^, and go awry in neurodegenerative disease^64–66^; and have propeled our ability to probe biochemical and biophysical function through the installation of chemical probes and payloads that cannot be genetically encoded^8,67,68^. The MccB/TeCH-tag system provides a new method for C-terminal protein activation that vastly expands the toolkit for protein semisynthesis and protein bioconjugation. MccB/TeCH-tag is broadly useful based on its ability to drive peptide bond-forming reactions to high yield by coupling them to ATP cleavage and can be integrated with existing protein chemistry technologies to fuel biological discovery.

## Acknowledgements

We thank S. Coyle, D. Sashital, T. Galateo, R. Rajasekaran, H. Bridge, W. Leiter, and members of the Weeks lab for helpful discussions. This work was supported in part by startup funds from the University of Wisconsin-Madison Department of Biochemistry and by an NIH Director’s New Innovator Award (DP2GM149548) to A.M.W. C.L.F. was supported in part by the UW-Madison Biotechnology Training Program under grant number NIH 5 T32 GM135066 and by a William H. Peterson Graduate Fellowship from the University of Wisconsin-Madison Department of Biochemistry.

## Author contributions

C.L.F, D.D. and A.M.W. designed the experiments. C.L.F. and D.D. performed experiments. C.L.F., D.D., and A.M.W. analyzed data. C.L.F. and A.M.W. wrote the manuscript.

## Methods

### Key chemicals and materials

Reagents screened for nucleophilic capture of MccB-activated C termini are listed in **Table S1**. Maleimide, Dibenzocyclooctyne-PEG4-biotin, and AlaPhe dipeptide were purchased from Sigma Aldrich. TCEP hydrochloride was purchased from Gold Biotechnology. Cyanine5 maleimide was purchased from Lumiprobe. EZ-Link-maleimide-PEG2-biotin was purchased from Thermo Fisher Scientific. Protected amino acids, 1-hydroxybenzotriazole, 4-alkoxybenzyl alcohol resins, and Rink amide resin for solid-phase peptide synthesis were purchased from Chem Impex International.

### Solid phase peptide synthesis

MccA C-terminal variant peptides with C-terminal carboxylate groups were synthesized using fluorenylmethyloxycarbonyl (Fmoc) chemistry on 4-alkoxy-benzyl-alcohol resin preloaded with the required C-terminal amino acid (Chem Impex International). All other peptides with C-terminal amides were synthesized using Rink Amide resin (Chem Impex International). Fmoc groups were deprotected via 30-minute incubation in 20% methylpiperidine in DMF (20% v/v). Coupling steps were performed with 5 molar equivalents of the appropriate Fmoc-protected amino acid, 5 molar equivalents of diisopropylcarboiimide (DIC), and 5 molar equivalents of 1-hydroxy-benzotriazole (HOBt). Completed peptides were cleaved from the resin via incubation in a cocktail containing 95% trifluoroacetic acid, 2.5% triisopropylsilane, and 2.5% water. Peptides were concentrated under a stream of nitrogen and precipitated with 10 volumes of diethyl ether. Precipitated peptides were washed with additional diethyl ether and allowed to dry. The resulting crude product was purified using an Agilent 1260 Infinity II HPLC fitted with a semi-preparative ZORBAX Eclipse XDB-C18, 9.4 × 250 mm, 5 μm column. Crude products were separated using a 30-minute gradient from 0 to 100% B (A=0.1% trifluoroacetic acid in water, B=acetonitrile). Selected fractions were lyophilized, resuspended in water, and stored at −20°C.

### Mass spectrometry

Electrospray ionization liquid chromatography mass spectrometry (ESI-LC-MS) analysis was performed on an Agilent 6230B time of flight (TOF) mass spectrometer. Samples containing peptide substrates were separated on a ZORBAX Eclipse XDB-C18, Solvent Saver Plus, 3 × 150 mm, 3.5 μm column using a 5-minute gradient from 0 to 100% B (A=0.1% formic acid in water, B=acetonitrile). Extracted ion chromatograms were generated using Agilent MassHunter Qualitative Analysis v10.0 and Agilent TOF Quantitative Analysis v11.0. Samples containing intact protein substrates were separated on a PLRP-S 1000 Å, 50×1 mm, 5 μm column using a 3.9-minute gradient from 20 to 60% B (A=0.1% formic acid in water, B=acetonitrile). The maximum entropy charge deconvolution algorithm in Agilent MassHunter BioConfirm v10.0 was used to determine the neutral mass of intact proteins. For mixtures of TeCH-tagged proteins, the pMod algorithm was applied for charge deconvolution in Agilent MassHunter BioConfirm v10.0

### Molecular biology and plasmid construction

Plasmids were constructed using standard Gibson assembly cloning methods with *E. coli* XL10 as the cloning host. Oligonucleotides were purchased from Integrated DNA Technologies, and plasmid sequences were confirmed via Sanger Sequencing performed by Quintara Biosciences or Functional Biosciences.

#### pBH4-MccB

A codon-optimized gene encoding *E. coli* MccB was purchased from Twist biosciences. The gene was inserted into the pBH4 vector between the BamHI and NotI restriction sites using Gibson assembly.

#### pBH4-eGFP-MccA fusions

eGFP was amplified from pBH4-eGFP (a gift from Scott Coyle). PCR products were inserted into pBH4 between BamHI and NotI restriction sites using Gibson assembly.

#### pBH4-Cysteine free-eGFP-MccA fusions

Cysteine-free eGFP was amplified from ss-cfSGFP2 (Addgene #37535). PCR products were inserted into pBH4 between BamHI and NotI restriction sites using Gibson assembly.

### Protein expression and purification

Protein expression and purification methods for each protein used in this study are described below. Following purification, the purity of proteins was analyzed by SDS-PAGE (**Fig. S19**).

#### MccB

##### E. coli MccB

MGHHHHHHDYDIPTTENLYFQGSMDYILGRYVKIARYGSGGLVGGGGKEQYVEDLALWENIIKTAYCFITPSSYTAALETVNIPEKDFSNCFRFLKENFFIIPSEYNNSTENNRYSRNFLHYQSYGANPVLVQDKLKDAKVVILGCGGIGNHVSVILATSGIGEIILIDNDQIENTNLTRQVLFSENDVGKNKTEVIKRELLKRNSEISVSEIALNINDYTDLHKVPEADIWVVSADHPFNLINWVNKYCVRANQPYINAGYVNDIAVFGPLYVPGKTGCYECQKVVADLYGSEKENIDHKIKLINSRFKPATFAPVNNVAAALCAADVIKFIGKYSEPLSLNKRIGIWSDEIKIHSQNMGRSPVCSVCGNRM

##### H. pylori MccB

MQWYQTSFSACVGQTDTENIIGLGTYQYCVDHNEFEKSLRLLVFLRMKKSMNEIKSFMETSKIEHNIFDKLVANKLITSFILNPNDKQNFKNHLFIDLVSNKPELTINNFKKTIFIIIGCGGIGNFVSYALASFYPKKLILLDKDTVDPSNLNRQFLFDKNYISQYKTSAIKQALSSRFGINIETVDDFASEDNLEEIFSKNKKENLFGIVSGDNPNTVQLATRFFCKCRIPFLNIGYLNDISLIGPFYIPSLSCCPFCHNSFALDDKKDGDKNLDIYLINDRMQAPSSFLNNSIASSLAISDIIQFMSNDFNSIKSLNCRFGVDNKTFKTYTLPSSVDYKCAFCSNHNL

##### L. johnsonii MccB

MFYKTSYLATGGCSNHQGILGVGTKQYFVSEADYLKSLKILDFLLNKKTYDEVIKFCEKNNINKSIFDTLVEHNLIVKENLYVEKKDDLNFKNKLYFHALGLNGNALAKEFADTTFVIVGCGGIGNFISFAIGSLSPRKIELIDGDKIEKSNLNRQFLFTENDIGKYKVDVLKKNLVERNNKLSISEYKEYVSKEVLHNIFEQNKKNKTLVILSGDSFSALSLTAKACVKSEIPFLNIGYLNDISAIGPFYIPGISSCPFCHNALSISDDISSGHNESKILEDRINANNEAPSSFTNNALAASMGIADIIEFLSHNYERINSLNKRFGINSATFEKYVLEVNRDRKCEICSHGE pBH4-MccB plasmids were transformed into chemically competent *E. coli* BL21(DE3) cells for overexpression. LB starter cultures (15 mL, 50 μg/mL carbenicillin) were grown overnight and used to inoculate 1 L LB cultures (50 μg/mL carbenicillin). Cultures were incubated at 37°C with vigorous shaking until OD_600_ reached ∼0.6. Cultures were chilled on ice for 15 minutes prior to adding 0.4 mM IPTG (isopropyl-β-D-thiogalactopyranoside). Cultures were shaken an additional 24 hours at 16°C. Cell pellets were harvested by centrifugation at 4°C, resuspended in 40 mL lysis buffer (25 mM Tris, 500 mM NaCl, 10 mM MgCl2, pH 8.0). Cells were lysed by three passes through an Emulsiflex microfluidizer at 15000 psi, and the resulting lysate was centrifuged at 8000 × *g* for 15 minutes at 4°C. Ni-NTA resin (1 mL) was added to the clarified lysate and His-tagged proteins were allowed to bind for 1 hour at 4°C with gentle rocking. The Ni-NTA resin was collected by centrifugation at 500 × *g* for 5 min, transferred to a 15 mL column, and washed with 15 mL wash buffer (20 mM Tris pH 8.0, 500 mM NaCl, 25 mM imidazole). Protein was eluted with 6 mL elution buffer (20 mM Tris pH 8.0, 500 mM NaCl, 200 mM imidazole) and dialyzed against wash buffer overnight at 4°C. The dialyzed protein was then concentrated using Amicon centrifugal filter units (10,000 MWCO) and further purified by size-exclusion chromatography (Superdex Hiload 75, GE Healthcare) using storage buffer (25 mM Tris pH 8.0, 50 mM NaCl, 1 mM DTT, 10% glycerol). Collected fractions were analyzed by Coomassie-stained SDS-PAGE. Fractions containing MccB were pooled, concentrated, and flash frozen in liquid nitrogen. Protein concentrations were determined using absorbance at 280 nm and the protein extinction coefficient as calculated using ProtParam.

#### MccB protein substrates

##### eGFP-MccA N7G

MGHHHHHHDYDIPTTENLYFQGSMVSKGEELFTGVVPILVELDGDVNGHKFSVSGEGEGDATYGKLTLKFICTTGKLPVPWPTLVTTLTYGVQCFSRYPDHMKQHDFFKSAMPEGYVQERTIFFKDDGNYKTRAEVKFEGDTLVNRIELKGIDFKEDGNILGHKLEYNYNSHNVYIMADKQKNGIKVNFKIRHNIEDGSVQLADHYQQNTPIGDGPVLLPDNHYLSTQSALSKDPNEKRDHMVLLEFVTAAGITL GMDELYK<u>MRTGNAG</u>

##### eGFP-1x-MccA N7G

MGHHHHHHDYDIPTTENLYFQGSMVSKGEELFTGVVPILVELDGDVNGHKFSVSGEGEGDATYGKLTLKFICTTGKLPVPWPTLVTTLTYGVQCFSRYPDHMKQHDFFKSAMPEGYVQERTIFFKDDGNYKTRAEVKFEGDTLVNRIELKGIDFKEDGNILGHKLEYNYNSHNVYIMADKQKNGIKVNFKIRHNIEDGSVQLADHYQQNTPIGDGPVLLPDNHYLSTQSALSKDPNEKRDHMVLLEFVTAAGITL GMDELYK<u>GGGGMRTGNAG</u>

##### Cysteine free eGFP-1x-N7G

MGHHHHHHDYDIPTTENLYFQGSMVSKGEELFTGVVPILVELDGDVNGHKFSVSGEGEGDATYGKLTLKFISTTGKLPVPWPTLVTTLTYGVQMFARYPDHMKQHDFFKSAMPEGYVQERTIFFKDDGNYKTRAEVKFEGDTLVNRIELKGIDFKEDGNILGHKLEYNYNSHNVYITADKQKNGIKANFKIRHNIEDGGVQLADHYQQNTPIGDGPVLLPDNHYLSTQSKLSKDPNEKRDHMVLLEFVTAAGITL GMDELYK<u>GGGGMRTGNAG</u>

##### MBP-1x-N7G

MGHHHHHHDYDIPTTENLYFQGSMKIEEGKLVIWINGDKGYNGLAEVGKKFEKDTGIKVTVEHPDKLEEKFPQVAATGDGPDIIFWAHDRFGGYAQSGLLAEITPDKAFQDKLYPFTWDAVRYNGKLIAYPIAVEALSLIYNKDLLPNPPKTWEEIPALDKELKAKGKSALMFNLQEPYFTWPLIAADGGYAFKYENGKYDIKDVGVDNAGAKAGLTFLVDLIKNKHMNADTDYSIAEAAFNKGETAMTINGPWAWSNIDTSKVNYGVTVLPTFKGQPSKPFVGVLSAGINAASPNKELAKEFLENYLLTDEGLEAVNKDKPLGAVALKSYEEELAKDPRIAATMENAQKGEIMPNIPQMSAFWYAVRTAVINAASGRQTVDEA LKDAQTNSSSNNNNNNNNNNLGIEGRG<u>GGGGMRTGNAG</u>

##### Protein L-1x-N7G

MHHHHHHKEETPETPETDSEEEVTIKANLIFANGSTQTAEFKGTFEKATSEAYAYADTLKKDNGEYTVDVADKGYTLNIKFAGKEKTPEEPKEEVTIKANLIYADGKTQTAEFKGTFEEATAEAYRYADALKKDNGEYTVDVADKGYTLNIKFAGKEKTPEEPKEEVTIKANLIYADGKTQTAEFKGTFEEATAEAYRYADLLAKENGKYTVDVADKGYTLNIKFAGKEKTPEEPKEEVTIKANLIYADGKTQTAEFKGTFAEATAEAYRYADLLAKENGKYTADLEDGGYTINI RFAGKKVDEKPEEKEQVTIKENIYFEDGTVQTATFKGTFAEATAEAYRYADLLSKEHGK YTADLEDGGYTINIRFA<u>GGGGSGGGSMRTGNAG</u>

##### PTP1B(1-321)-1x-N7G

MGHHHHHHDYDIPTTENLYFQGSMEMEKEFEQIDKSGSWAAIYQDIRHEASDFPCRVAKLPKNKNRNRYRDVSPFDHSRIKLHQEDNDYINASLIKMEEAQRSYILTQGPLPNTCGHFWEMVWEQKSRGVVMLNRVMEKGSLKCAQYWPQKEEKEMIFEDTNLKLTLISEDIKSYYTVRQLELENLTTQETREILHFHYTTWPDFGVPESPASFLNFLFKVRESGSLSPEHGPVVVHCSAGIGRSGTFCLADTCLLLMDKRKDPSSVDIKKVLLEMRKFRMGLIQTADQLRFSYLAVIEGAKFIMGDSSVQDQWKELS HEDLEPPPEHIPPPPRPPKRILEPHN<u>GGGGMRTGNAG</u>

The appropriate plasmid was transformed into chemically competent *E. coli* BL21(DE3) cells for overexpression. LB starter cultures (15 mL, 50 μg/mL carbenicillin) were grown overnight and used to inoculate 1 L LB cultures (50 μg/mL carbenicillin). Cultures were incubated at 37°C with vigorous shaking until OD_600_ reached ∼0.6. Cultures were chilled on ice for 15 minutes prior to adding 0.25 mM IPTG (isopropyl-β-D-thiogalactopyranoside). Cultures were shaken an additional 16 hours at 18°C. Cell pellets were harvested by centrifugation at 4°C, resuspended in 40 mL wash buffer (50 mM sodium phosphate, 300 mM NaCl, 20 mM imidazole, pH 8.0). Cells were lysed by three passes through an Emulsiflex microfluidizer at 15000 psi, and the resulting lysate was centrifuged at 8000 × *g* for 15 minutes at 4°C. Ni-NTA resin was added to the clarified lysate and allowed to bind for 1 hour at 4°C with gentle rocking. The Ni-NTA resin was collected by centrifugation at 500 × *g* for 5 min, transferred to a 15 mL column, and washed with 15 mL wash buffer. Protein was eluted with 6 mL elution buffer (50 mM sodium phosphate, 300 mM NaCl, 250 mM imidazole, pH 8.0) and dialyzed overnight at 4°C against wash buffer containing 1 mM DTT. To remove the His tag, TEV protease was added to the dialysis tubing at a ratio of 1:50 protease:substrate. After 16-24 hours digestion, the protein solution was passed through 2 mL equilibrated Ni-NTA resin. The resulting protein was then concentrated using Amicon centrifugal filter units (10,000 MWCO) and further purified by size-exclusion chromatography (Superdex Hiload 75, GE Healthcare) using storage buffer (20 mM Tris pH 8.0, 150 mM NaCl, 10% glycerol). Collected fractions were analyzed by Coomassie-stained SDS-PAGE. Fractions containing the correct protein molecular weight were pooled, concentrated, and flash frozen in liquid nitrogen. Protein concentrations were determined using absorbance at 280 nm and the protein extinction coefficient as calculated using ProtParam.

#### Anti-eGFP fab Heavy Chain-1x-N7G

##### Light chain

MKSLLPTAAAGLLLLAAQPAMASDIQMTQSPSSLSASVGDRVTITCRASQSVSSAVAWYQQKPGKAPKLLIYSASSLYSGVPSRFSGSRSGTDFTLTISSLQPEDFATYYCQQSWGLITFGQGTKVEIKRTVAAPSVFIFPPSDSQLKSGTASVVCLLNNFYPREAKVQWKVDNALQSGNSQESVTEQDSKDSTYSLSSTLTLSKADYEKHKVYACEVTHQGLSSPVTKSFNRGEC

##### Heavy chain

MKKNIAFLLASMFVFSIATNAYAEISEVQLVESGGGLVQPGGSLRLSCAASGFNISYYSIHWVRQAPGKGLEWVASIYPYYSSTSYADSVKGRFTISADTSKNTAYLQMNSLRAEDTAVYYCARAGWVASSGMDYWGQGTLVTVSSASTKGPSVFPLAPSSKSTSGGTAALGCLVKDYFPEPVTVSWNSGALTSGVHTFPAVLQSSGLYSLSSVVTVPSSSLGTQTYICNVNHKPSNTKVDKKVEPKSCGGGG GMRTGNAG

pPSL937-anti-GFP^69^ Fab-heavychain-1xN7G was transformed into chemically competent C43 Pro+ pTUM protease-deficient *E. coli* cells for overexpression. LB starter cultures (5 mL, 50 μg/mL carbenicillin, 25 μg/mL chloramphenicol) were grown overnight and used to inoculate 1 L TB expression cultures (50 μg/mL carbenicillin, 25 μg/mL chloramphenicol, 0.05% w/v glucose, 0.5% w/v lactose, 1% w/v galactose, 2 mM MgSO_4_). Cultures were incubated at 37°C for 6 hours and grown for another 18 hours at 30°C. Cell pellets were harvested by centrifugation at 4°C and resuspended in 40 mL 1:1 PBS:Bacterial Protein Extraction Reagent (BPER, ThermoFisher Scientific). Cell pellets were lysed by incubating in 60°C water bath for 30 minutes. The resulting lysate was centrifuged at 8000 × *g* for 15 minutes at 4°C. The supernatant was filtered with 0.2 μm syringe filters and loaded onto a HiTrap Protein A HP column (Cytiva). The column was washed with PBS and protein was eluted with 100 mM acetic acid and immediately neutralized. Fractions were analyzed by Coomassie-stained SDS-PAGE. Fractions containing the correct protein molecular weight were pooled, concentrated, and flash frozen in liquid nitrogen. Protein concentrations were determined using absorbance at 280 nm and the protein extinction coefficient as calculated using ProtParam.

#### Subtiligase

AQSVPYGVSQIKAPALHSQGYTGSNVKVAVIDSGIDSSHPDLKVAGGASFVPSETNPFQDNNSHGTHVAGTVAALDNSIGVLGVAPSASLYAVKVLGADGSGQYSWIISGIEWAIANNMDVINLALGGPSGSAALKAAVDKAVASGVVVVAAAGNEGTSGSSSTVGYPGKYPSVIAVGAVDSSNQRASFSSVGPELDVMAPGVSIQSTLPGNRYGAYSGTCMASAHVAGAAALILSKHPNWTNTQVRSSLENTTTKLGDSFYYGKGLINVQAAAQ

pBS42-pre-pro-degra subtiligase-His6 (M50F, N76D, N109S, M124L, S125A, K213R, N218S) was transformed into *B. subtilis* BG2864 for expression and secretion. 2xYT starter cultures (15 mL, 12.5 μg/mL chloramphenicol) were grown overnight and used to inoculate 200 mL 2xYT cultures (12.5 μg/mL chloramphenicol, 5 mM CaCl_2_) to OD_600_ 0.03-0.05. Cultures were incubated at 37°C with vigorous shaking for 24 hours. Cells were pelleted by centrifugation at 4°C, and the resulting supernatant was added to 3 volumes of ice-cold ethanol. The precipitate was harvested by centrifugation at 4°C and resuspended in wash buffer (50 mM sodium phosphate, 300 mM NaCl, 20 mM imidazole, pH 8.0). After brief centrifugation, Ni-NTA resin (400 μL) was added to the resuspended pellet and His-tagged protein was allowed to bind for 1 hour at 4°C with gentle rocking. The Ni-NTA resin was collected by centrifugation at 500 × *g* for 5 min, transferred to a 1 mL spin column, and washed with 4 mL wash buffer. Protein was eluted with 0.8 mL elution buffer (50 mM sodium phosphate, 300 mM NaCl, 250 mM imidazole, pH 8.0). The resulting protein was then concentrated using Amicon centrifugal filter units (10,000 MWCO) and further purified by size-exclusion chromatography (Superdex Hiload 75, GE Healthcare) using storage buffer (100 mM bicine, 5 mM DTT, 10% glycerol, pH 8.5). Collected fractions were analyzed by Coomassie-stained SDS-PAGE. Fractions containing the correct protein molecular weight were pooled, concentrated, and flash frozen in liquid nitrogen. Protein concentrations were determined using absorbance at 280 nm and the protein extinction coefficient as calculated using ProtParam.

### Sequence similarity network construction

A sequence similarity network (SSN) for representative members of the E1-like/ThiF superfamily was constructed using the Enzyme Function Initiative Enzyme Similarity Tool^70,71^ at https://efi.igb.illinois.edu/efi-est/. Input sequences consisted of those retrieved from the ThiF family in Pfam^72^ and were comprised of reviewed Uniprot^73^ entries and MccB sequences with annotated MccA substrates from a bioinformatic analysis of MccB homologs involved in biosynthesis of microcin C. The SSN was visualized in Cytoscape 3.9.1 using a minimum alignment score of 100.

### Purine nucleoside phosphatase (PNP) kinetic assays

The kinetics of MccB-catalyzed adenylation were characterized using a previously reported coupled PNP enzyme assay and reagents supplied in the EnzCheck Pyrophosphate Assay Kit (Thermo Fisher Scientific). To screen MccB for adenylation activity with C-terminal variants of MccA, 5 μM of the appropriate MccB variant was incubated with 0.25 mM of each MccA variant in a reaction with 0.25 mM ATP, 5 mM MgCl_2_, 0.5 U/mL purine nucleoside phosphorylase (PNP), and 0.5 U/mL inorganic pyrophosphatase (IP). To screen *E. coli, H. pylori,* and *L. johnsonii* MccB homologs for adenylation activity with MccA homologs, 5 μM of the appropriate MccB enzyme was incubated with 0.25 mM of each MccA variant in a reaction with 0.25 mM ATP, 5 mM MgCl2, 0.5 U/mL purine nucleoside phosphorylase (PNP), and 0.5 U/mL inorganic pyrophosphatase (IP). Reactions containing MccA-N7G peptides also contained 5 mM Mesna and 5 mM TCEP. To collect kinetic parameters for individual peptide substrates, the peptide concentration was varied from 0-600 μM in a reaction with 5 μM MccB, 0.25 mM ATP, 5 mM MgCl_2_, 0.5 U/mL purine nucleoside phosphorylase (PNP), and 0.5 U/mL inorganic pyrophosphatase (IP). Reactions with MccA-N7G were performed using 0.5 μM MccB and peptide concentrations from 0-100 μM). Reactions were initiated with the addition of MccB and absorbance at 360 nm was monitored using a Tecan plate reader. Initial rates of pyrophosphate production were calculated based on the initial rate of absorbance change and the resulting purine analog extinction coefficient (11,000 M^−1^cm^−1^). Reactions were carried out in triplicate and data analysis was performed using GraphPad Prism 10.

### LC-TOF MS analysis of MccB reactions with synthetic peptide substrates

For analysis of formation of *N*-AMPylated MccA, MccA substrate (250 μM) was incubated with 5 μM MccB and 5 mM ATP in reaction buffer (75 mM Tris pH 8.0, 5 mM MgCl_2_). Reactions were incubated for 16 hours at room temperature and quenched with addition of an equivalent volume of 0.6% TFA. These reaction conditions were used for MccBs from *E. coli*, *H. pylori*, and *L. johnsonii*. Quenched reactions were centrifuged at 8000 × *g* for 10 minutes and analyzed using LC-TOF MS as described in **Mass spectrometry**.

For analysis of *E. coli* MccB-catalyzed thioesterification, MccA-N7G substrate (250 μM) was incubated with 5 μM MccB, 5 mM ATP, 5 mM Mesna, and 5 mM TCEP in reaction buffer (75 mM Tris pH 8.0, 5 mM MgCl_2_). Reactions were incubated for 16 hours at room temperature and quenched with addition of an equivalent volume of 0.6% TFA. To screen the ability of *E. coli* MccB, *H. pylori* MccB, *L. johnsonii* MccB to catalyze thioesterification of MccA-N7Gs derived from each species, reactions contained 5 μM MccB, 5 mM ATP, 50 mM Mesna, and 12.5 mM TCEP in reaction buffer (75 mM Tris pH 8.0, 5 mM MgCl_2_). Quenched reactions were centrifuged at 8000 × *g* for 10 minutes and analyzed using LC-TOF MS as described in **Mass spectrometry**.

### Screening nucleophiles for MccB-mediated ligation of synthetic peptide substrates

Nucleophile solutions were prepared as aqueous solutions and adjusted to neutral pH using paper pH strips. Solutions were prepared in bulk and stored at −80°C until further use, unless otherwise noted. 250 μM MccA N7G was incubated with 5 μM MccB, 5 mM ATP, and the corresponding concentration of nucleophile in reaction buffer (25 mM Tris pH 8.0, 10 mM MgCl2). Reactions were incubated for 16 hours at room temperature and quenched with addition of an equivalent volume of 0.6% TFA. Quenched reactions were centrifuged at 8000 × *g* for 10 minutes and analyzed using LC-TOF MS as described in **Mass spectrometry**. Following identity assignment of peaks, percent conversion of the ligation product was calculated using the integrated peak areas of reactant peptide and product. Extracted ion chromatogram peak areas were calculated using Agilent TOF Quantitative Analysis v11.0.

### MccB-mediated C-terminal ligation of MccA-N7G with Cys

MccA-N7G (250 μM) was incubated for 16 hours at room temperature in reactions containing 5 μM MccB, 5 mM ATP, and the appropriate concentration of Cys/TCEP in reaction buffer (25 mM Tris pH 8.0, 10 mM MgCl_2_). For maleimide functionalization, reactions were performed with 1 mM Cys/TCEP for 16 hours followed by a 1-hour incubation with 1.5 mM maleimide. All reactions were quenched, centrifuged at 8000 × *g* for 10 minutes, and analyzed using LC-TOF MS as described in **Mass spectrometry**.

### MccA-N7G thioester exchange with N-terminal Cys peptide

C-terminal thioester was prepared in reactions containing 250 μM MccA-N7G, 5 μM MccB, 5 mM ATP, and 5 mM Mesna/TCEP in reaction buffer (25 mM Tris pH 8.0, 10 mM MgCl_2_). After incubating for 16 hours at room temperature, 20 mM CysTrp peptide was added to each reaction and incubated for an additional 4 hours. Reactions were quenched with the addition of an equivalent volume of 0.6% TFA. Quenched reactions were centrifuged at 8000 × *g* for 10 minutes and analyzed using LC-TOF MS as described in **Mass spectrometry**.

### MccB-mediated ligation of eGFP-MccA N7G protein substrates

Nucleophile solutions were prepared as aqueous solutions and adjusted to neutral pH using paper pH strips. Solutions were prepared in bulk and stored at −80°C until further use, unless otherwise noted. GFP-MccA-N7G variant (50 μM) was incubated with 5 μM MccB, 5 mM ATP, and the appropriate concentration of cysteine or Mesna/TCEP in reaction buffer (25 mM Tris pH 8.0, 10 mM MgCl_2_). Reactions were incubated at room temperature for the indicated times. For labeling time course experiments, reactions were quenched with the addition of an equivalent volume of 0.6% TFA. For maleimide functionalization of Cys-ligated proteins, reactions were desalted using 75 μL 7K MWCO Zeba Micro Spin Desalting Columns (Thermo Fisher Scientific) and incubated with maleimides overnight at 4°C. Reactions were centrifuged at 8000 × *g* for 10 minutes and analyzed using LC-TOF MS as described in **Mass spectrometry**.

### GFP-MccA N7G thioester exchange time course with CysTrp dipeptide

C-terminal thioester was prepared in reactions containing 50 μM cysteine-free GFP-1x-N7G, 5 μM MccB, 5 mM ATP, and 5 mM Mesna/TCEP in reaction buffer (25 mM Tris pH 8.0, 10 mM MgCl_2_). After incubating for 1 h at room temperature reactions were desalted using 75 μL 7K MWCO Zeba Micro Spin Desalting Columns (Thermo Fisher Scientific). The peptide was added for a final concentration of 5 mM. Reactions were quenched at various time points by addition of an equivalent volume of 0.6% TFA. Quenched reactions were centrifuged at 8000 × *g* for 10 minutes and analyzed using LC-TOF MS as described in **Mass spectrometry**.

### GFP-MccA N7G expressed protein ligation with N-terminal Cys peptide followed by copper-free click chemistry

Reactions contained 50 μM cysteine free eGFP-1x-N7G, 5 μM MccB, 5 mM ATP, 5 mM MESNA/DTT, and 5 mM CGAGS-azidoAla peptide in reaction buffer (25 mM Tris pH 8.0, 10 mM MgCl_2_). After incubating for 4 h at room temperature, reactions were desalted using 75 μL 7K MWCO Zeba Micro Spin Desalting Columns (Thermo Fisher Scientific). Desalting was repeated twice more. Dibenzocyclooctyne-PEG4-biotin was added to a final concentration of 3 mM and incubated for 1 h at room temperature. Reactions were desalted once more and DTT added to a final concentration of 5 mM for a 30-minute incubation at room temperature. Reactions were centrifuged at 8000 × *g* for 10 minutes and analyzed using LC-TOF MS as described in **Mass spectrometry**.

### MccB/subtiligase-catalyzed ligation of protein substrates

MccB/subtiligase-catalyzed C-terminal amide bond formation was performed in one-pot reactions containing 50 μM TeCH-tagged protein, 5 μM MccB, 5 mM ATP, 5 μM subtiligase, 5 mM peptide substrate, and 5 mM Mesna/DTT in reaction buffer (25 mM Tris pH 8.0, 10 mM MgCl_2_). After incubating for appropriate time at room temperature, reactions were diluted in 75 mM HEPES pH 8.0 and analyzed using LC-TOF MS as described in **Mass spectrometry**.

### MccB/subtiligase-mediated ligation of anti-GFP rAb followed by copper free click chemistry

C-terminal bioconjugation was performed in one-pot reactions containing 50 μM protein, 5 μM MccB, 5 mM ATP, 5 μM subtiligase, 5 mM peptide substrate, and 5 mM Mesna in reaction buffer (75 mM Tris pH 8.0, 5 mM MgCl_2_). After incubating for the appropriate time at room temperature, reactions were twice buffer exchanged into 75 mM HEPES, pH 8.0 using 500 μL 7K MWCO Zeba Micro Spin Desalting Columns (Thermo Fisher Scientific), and analyzed using LC-TOF MS. Absorbance at 280 nm was used to estimate the resulting protein concentration, and 2-5 molar equivalents of the DBCO reagent in DMSO were added to each sample. After 4 hours at room temperature, the samples were twice buffer exchanged as before, and analyzed using LC-TOF MS as described in **Mass spectrometry**.

### Cell culture and immunofluorescence

HEK293T-derived cells were grown in DMEM supplemented with 10% fetal bovine serum, 100 U/mL penicillin, 100 μg/mL streptomycin, and other antibiotics as appropriate. Cells were tested every six months for mycoplasma contamination using the LookOut Mycoplasma PCR Detection Kit (Sigma-Aldrich) according to the manufacturer’s instructions. Flp-In T-Rex 293 cells (ThermoFisher Scientific) were modified by introduction of doxycycline-inducible GFP targeted to the cell surface according to manufacturer’s instructions. To target GFP to the cell surface, an N-terminal Igκ signal peptide and a C-terminal PDGF receptor β-chain transmembrane domain were added to GFP to construct the sequence given below. Cells were seeded at 10,000 cells per well in a 96-well plate and grown to 50% confluency. Doxycycline (1 ug/mL) was then added to induce cell surface GFP expression. After 20 hours, cells were washed three times with ice cold PBS, fixed with PBS + 4% paraformaldehyde for 10 minutes, and washed three times with PBS + 3% BSA. Cells were stained with 10 μg/mL αGFP-rAb in PBS + 3% BSA for 1 hour at room temperature and washed three times with PBS + 3% BSA. Imaging was performed on an Echo Revolve epifluorescence microscope in the inverted configuration.

### Igκ-GFP-PDGFR-TM sequence

The N-terminal Igκ leader sequence is underlined, a V5 tag is shown in italic, and the C-terminal PDGF receptor transmembrane domain is underlined.

<u>METDTLLLWVLLLWVPGSTG</u>*GKPIPNPLLGLDSTGS*GGGASMVSKGEELFTGVVPILVELDGD VNGHKFSVSGEGEGDATYGKLTLKFICTTGKLPVPWPTLVTTLTYGVQCFSRYPDHMKQHDFFKSAMPEGYVQERTIFFKDDGNYKTRAEVKFEGDTLVNRIELKGIDFKEDGNILGHKLEYNYNSHNVYIMADKQKNGIKVNFKIRHNIEDGSVQLADHYQQNTPIGDGPVLLPDNHYLSTQSALSKDPN EKRDHMVLLEFVTAAGITLGMDELYKGSGGSGGGGS<u>AVGQDTQEVIVVPHSLPFKVVVISAILA</u> <u>LVVLTIISLIILIMLWQKKPR</u>

### LC-TOF MS analysis of MccB-catalyzed *N*-AMPylation of MccA positional scanning peptide library

MccA variants were synthesized individually in-house (according to **Solid-phase peptide synthesis**) or were purchased from Peptide2.0. For analysis of formation of *N*-AMPylated MccA variant, MccA substrate (250 μM) was incubated with 5 μM MccB and 5 mM ATP in reaction buffer (75 mM Tris pH 8.0, 5 mM MgCl_2_). Reactions were incubated for 16 hours at room temperature and quenched with addition of an equivalent volume of 0.6% TFA. Quenched reactions were centrifuged at 8000 × *g* for 10 minutes and analyzed using LC-TOF MS as described in **Mass spectrometry**.

### MccB homolog reactions with GFP-TeCH-tag fusion proteins

*E. coli, H. pylori,* and *L. johnsonii* were screened for protein thioesterification using eGFP fused with the MccA-N7G homolog sequences. Reactions contained 5 μM MccB, 5 mM ATP, 50 μM eGFP-TeCH fusion, 50 mM Mesna, and 12.5 mM TCEP in reaction buffer (75 mM Tris-HCl pH 8.0, 5 mM MgCl_2_). For multiplexed reactions containing all three eGFP-TeCH fusions, each eGFP-TeCH fusion was added to a final concentration of 16 μM. Reactions were quenched after 16 hours at room temperature by adding two volumes of 0.6% TFA. Samples were analyzed using LC-TOF MS.

