## Supplemental Information for "Engineered reactivity of a bacterial E1-like enzyme enables ATP-driven modification of protein C termini"

**Table S1. Chemical reagents used for nucleophilic capture of MccB-activated C-termini.**

| Reagent | CAS No. | supplier |
|---|---|---|
| hydroxylamine hydrochloride | 5470-11-1 | Sigma Aldrich |
| O-methylhydroxylamine hydrochloride | 593-56-6 | Sigma Aldrich |
| O-ethylhydroxylamine hydrochloride | 3332-29-4 | Sigma Aldrich |
| O-(Prop-2-yn-1-yl)hydroxylamine hydrochloride | 21663-79-6 | AmBeed |
| hydrazine monohydrate | 7803-57-8 | Sigma Aldrich |
| Prop-2-yn-1-ylhydrazine hydrochloride | 1187368-95-1 | Enamine |
| N-(But-3-yn-1-yl)-4-hydrazineylbenzamide | 2081100-00-5 | Sigma Aldrich |
| propylamine hydrochloride | 556-53-6 | Sigma Aldrich |
| butylamine | 109-73-9 | Sigma Aldrich |
| tert-butylamine hydrochloride | 10017-37-5 | Sigma Aldrich |
| benzylamine hydrochloride | 3287-99-8 | Sigma Aldrich |
| cyclohexylamine | 108-91-8 | Sigma Aldrich |
| allylamine hydrochloride | 10017-11-5 | Fisher Scientific |
| propargylamine | 2450-71-7 | AK Scientific |
| dimethylamine hydrochloride | 506-59-2 | Sigma Aldrich |
| N-acetyl cysteine | 616-91-1 | Sigma Aldrich |
| N-acetyl cysteamine | 1190-73-4 | Sigma Aldrich |
| dithiothreitol | 27565-41-9 | Gold Biotechnology |
| Mesna | 19767-45-4 | Selleckchem |
| 4-mercaptophenylacetic acid | 39161-84-7 | Fisher Scientific |
| Cysteine | 52-90-4 | Sigma Aldrich |

**Figure S1. C-terminal MccA variants screened by continuous coupled enzyme assay for pryophosphate release.** (A) Rate of pyrophosphate release by MccB in presence of 250 μM of the corresponding C-terminal MccA variant. Absorbance is converted to molarity using the MESG extinction coefficient. (B) Raw absorbance values used to calculate rates.

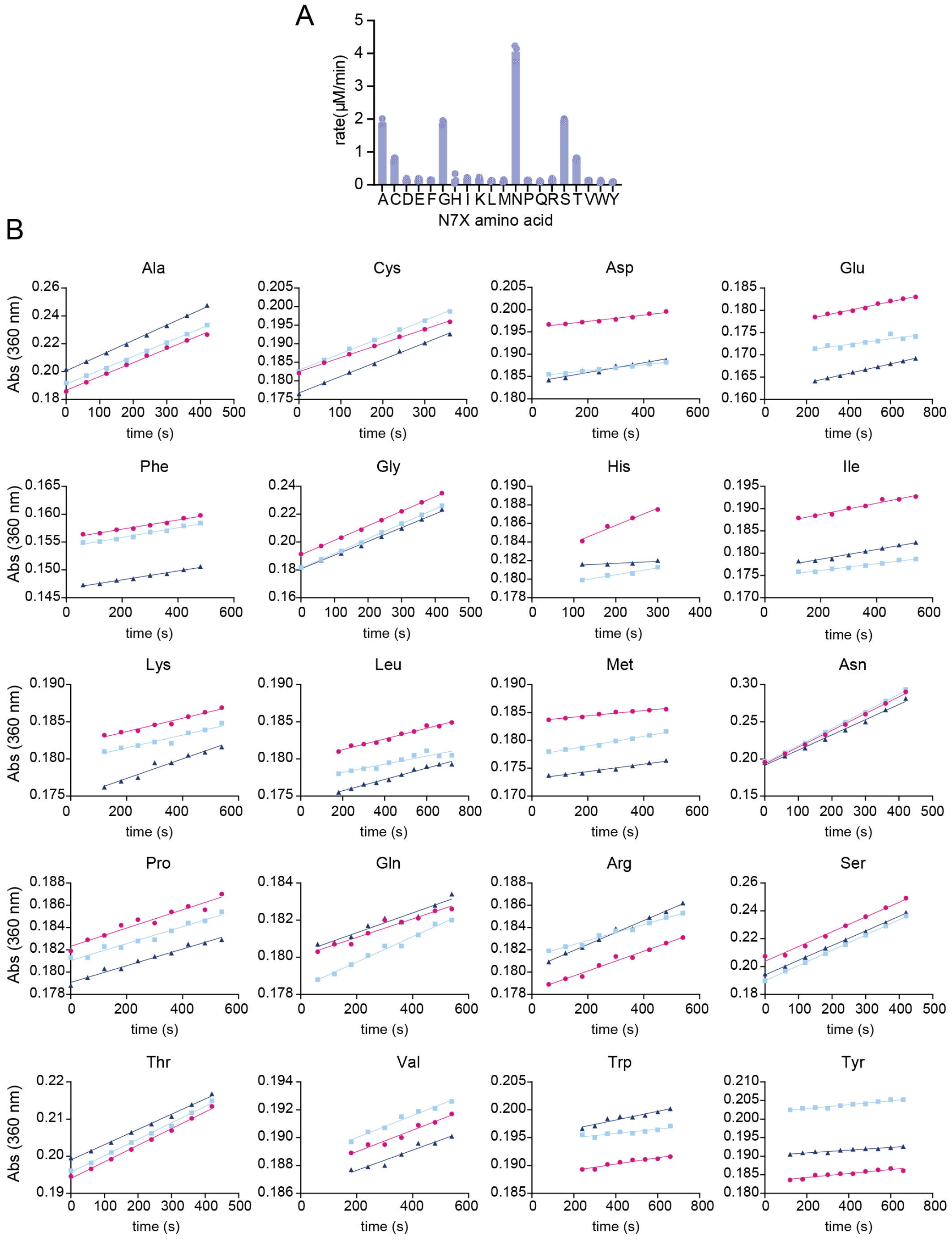

**Figure S2. Michaelis-Menten analysis of MccB-catalyzed pyrophosphate release using MccA C-terminal variants as substrates.** Assays were performed using the MccB concentration indicated on each plot. (A) Wild-type MccA. (B) MccA-N7A. (C) MccA-N7G. (D) MccA-N7S. (E) MccA-N7T.

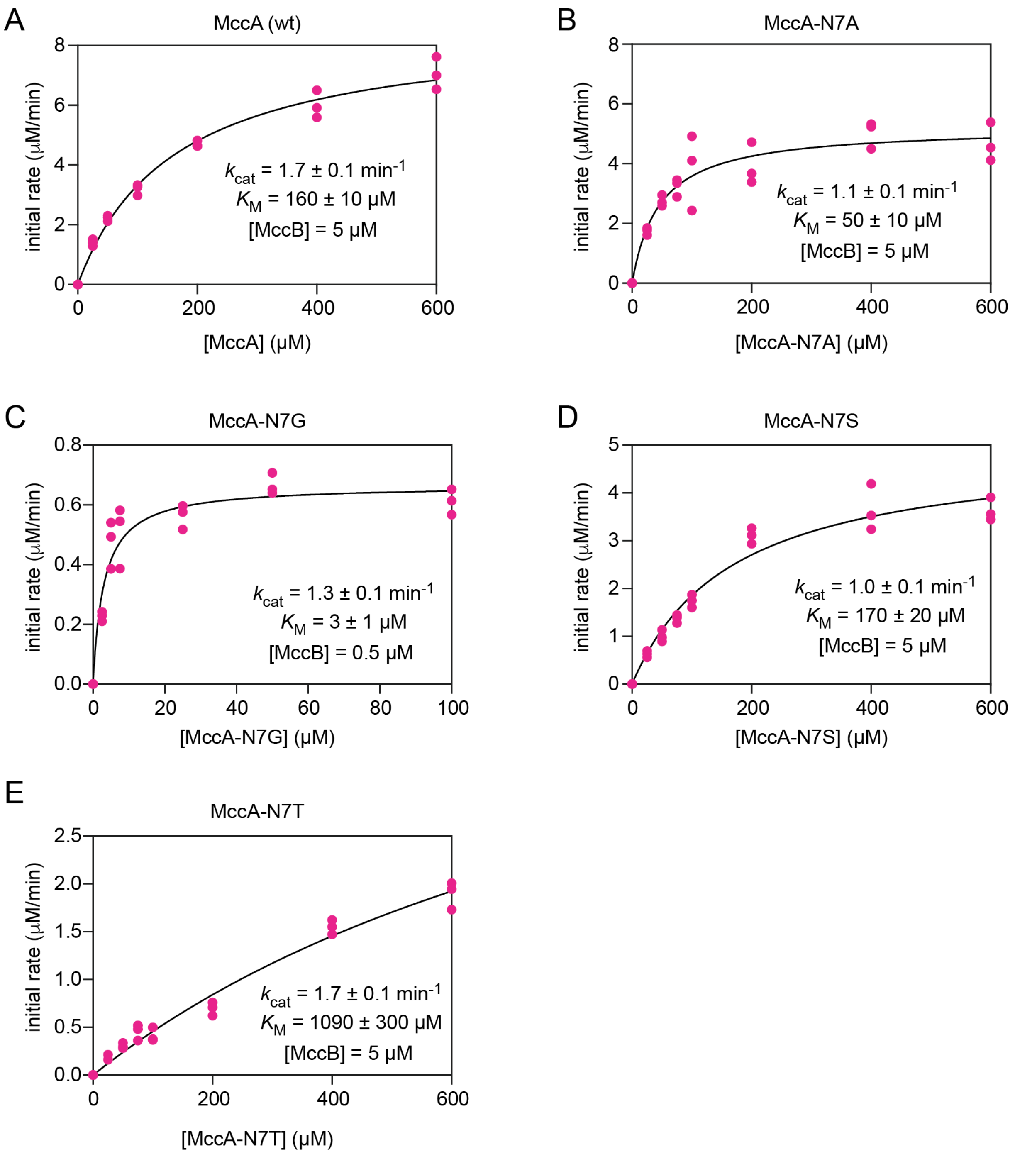

**Figure S3. C-terminal MccA variants screened by HPLC-MS for formation of N-AMPylated product.** (A) Percent product generated by MccB in presence of 250 μM of the corresponding C-terminal MccA variant. (B) Extracted ion chromatograms for C-terminal variant reactant and N-AMPylated product.

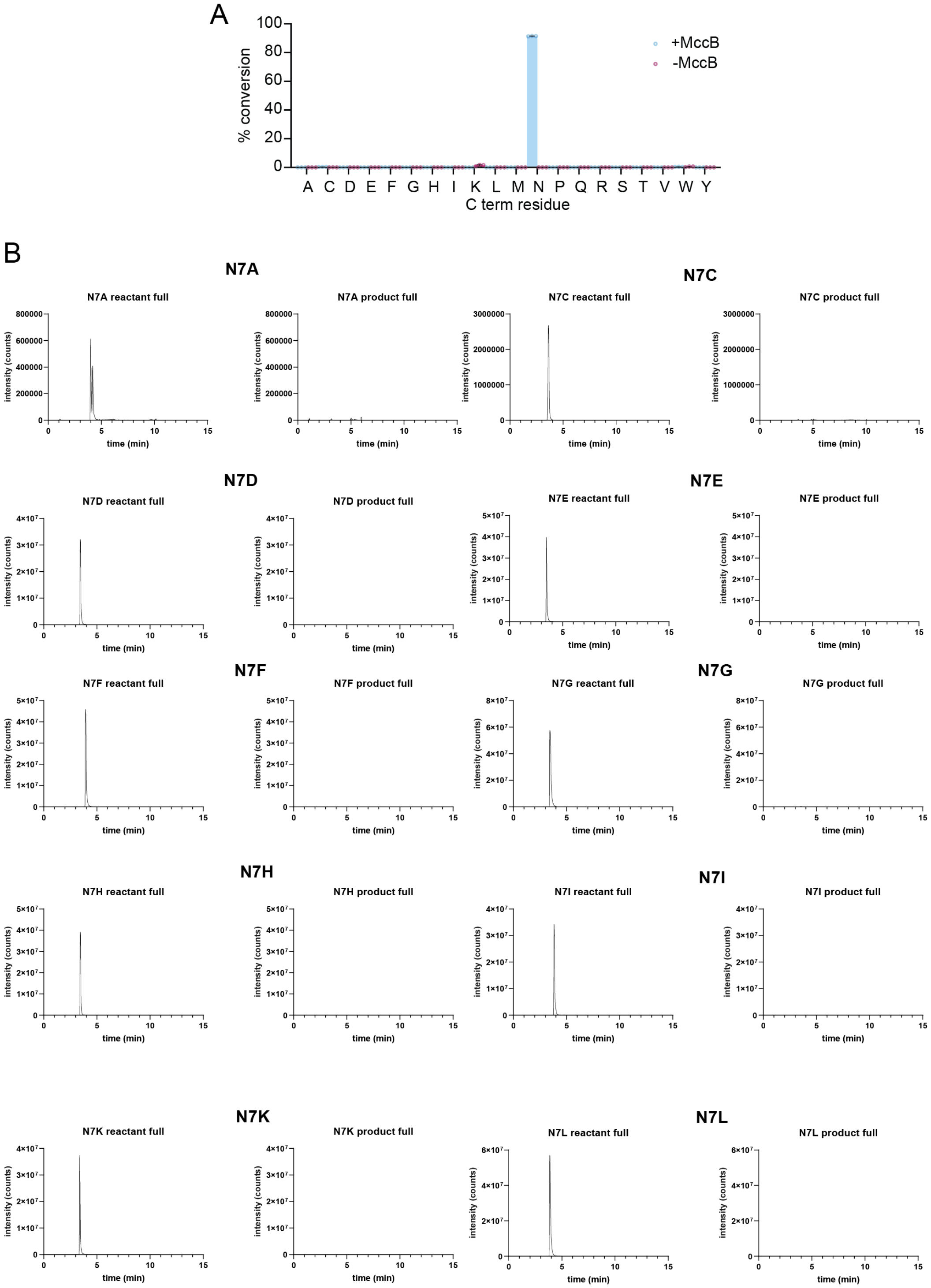

**Figure S3 (continued).**

B

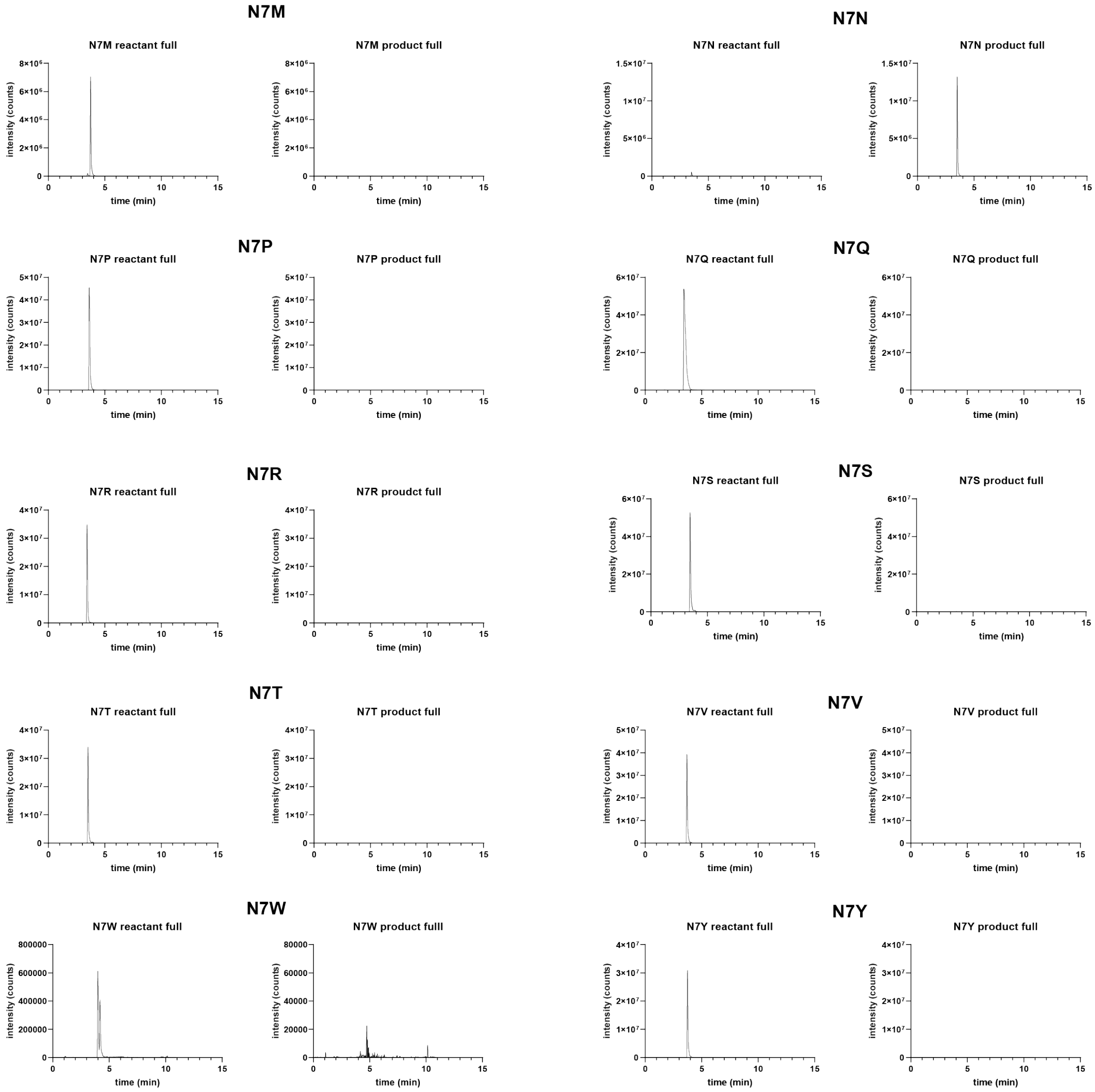

**Figure S4. HPLC-MS quantification of nucleophile ligation of MccA-N7G.** (A) Quantification of percent modified MccA-N7G generated by MccB in presence of 150 mM nitrogen nucleophile, 250 μM peptide substrate, and 5 mM ATP after a 16 h reaction. (B) Quantification of percent modified MccA-N7G generated by MccB in presence of 150 mM thiol nucleophile, 250 μM peptide substrate, and 5 mM ATP after a 16 h reaction.

A

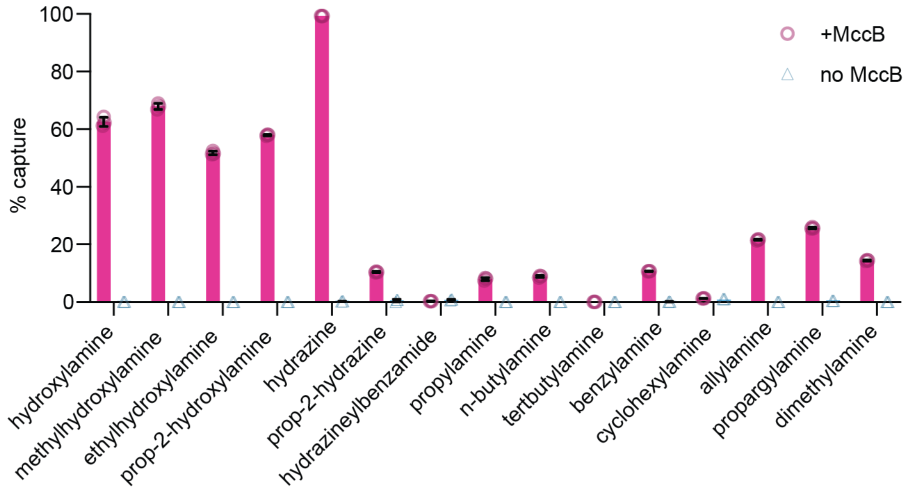


B

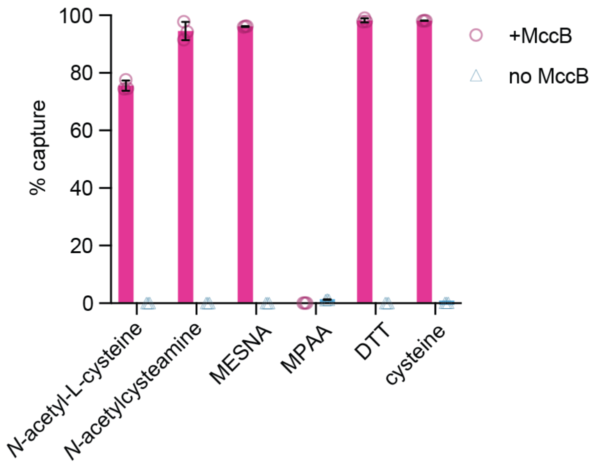

**Figure S5. Transthiolation of MccA N7G Mesna thioester with dithiothreitol (DTT).** (A) Scheme of initial Mesna thioesterification of MccA N7G followed by transthiolation with competing DTT. (B) Extracted ion chromatograms of the MccA N7G reactant, Mesna thioester product, and DTT thioester product. The masses of both singly and doubly charged species are included. (C) HPLC-MS quantification of DTT transthiolation of MccA N7G thioester.

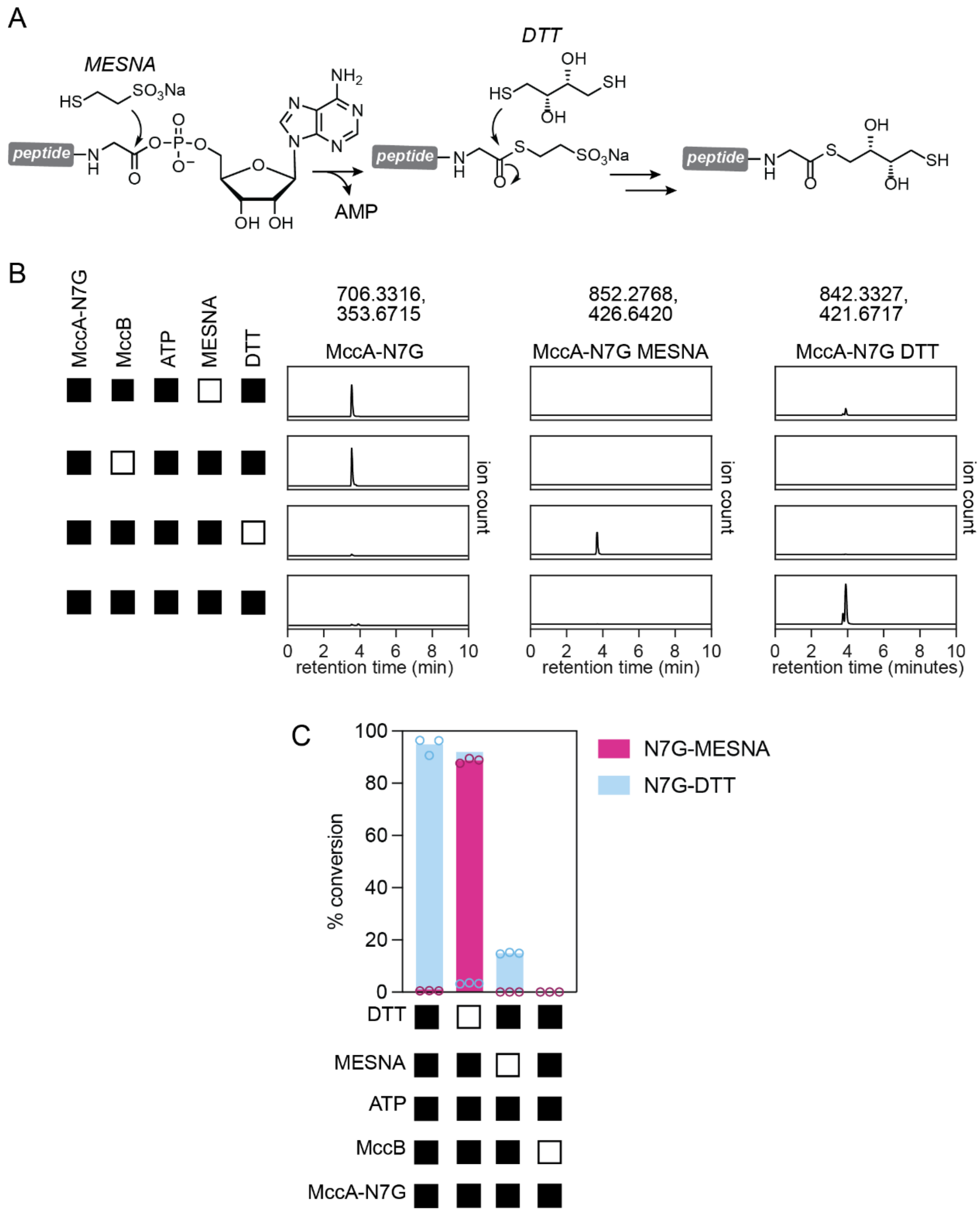

**Figure S6. Transthiolation of MccA N7G Mesna thioester with 4-mercaptophenylacetic acid (MPAA).** (A) Scheme of initial Mesna thioesterification of MccA N7G followed by transthiolation with competing MPAA. (B) Extracted ion chromatograms of the MccA N7G reactant, Mesna thioester product, and MPAA thioester product. The masses of both singly and doubly charged species are included. (C) HPLC-MS quantification of MPAA transthiolation of MccA N7G thioester.

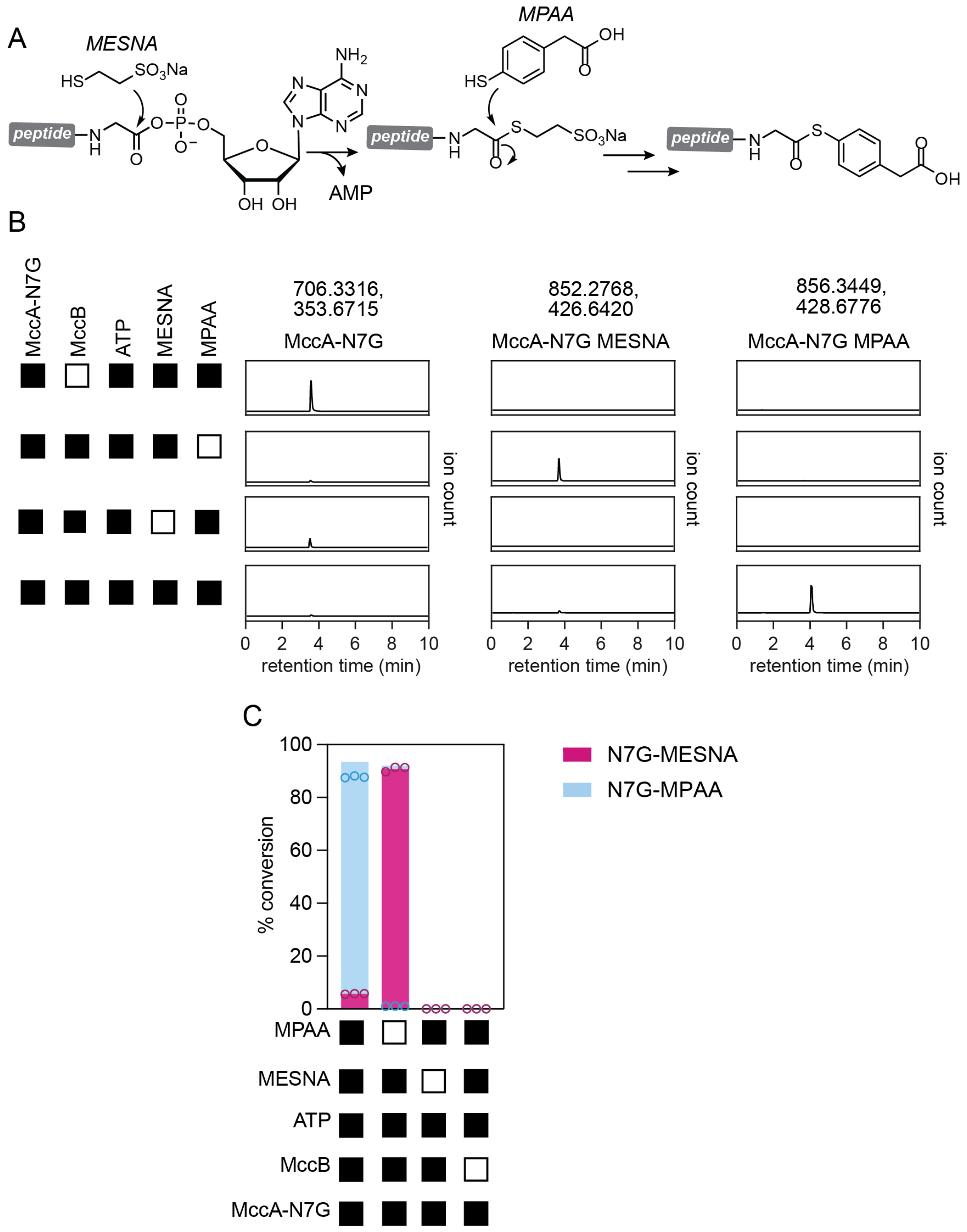

**Figure S7. Cysteine thioesterification of MccA-N7G undergoes *S*-to-*N* acyl shift to form a new amide bond.** (A) Scheme of initial Cysteine thioesterification of MccA-N7G followed by S-to-N acyl shift to free the thiol side chain for maleimide ligation. (B) Extracted ion chromatograms of the MccA N7G reactant, Cys captured product, and Cys-maleimide product. The masses of both singly and doubly charged species are included. (C) HPLC-MS quantification of Cys capture and maleimide modification.

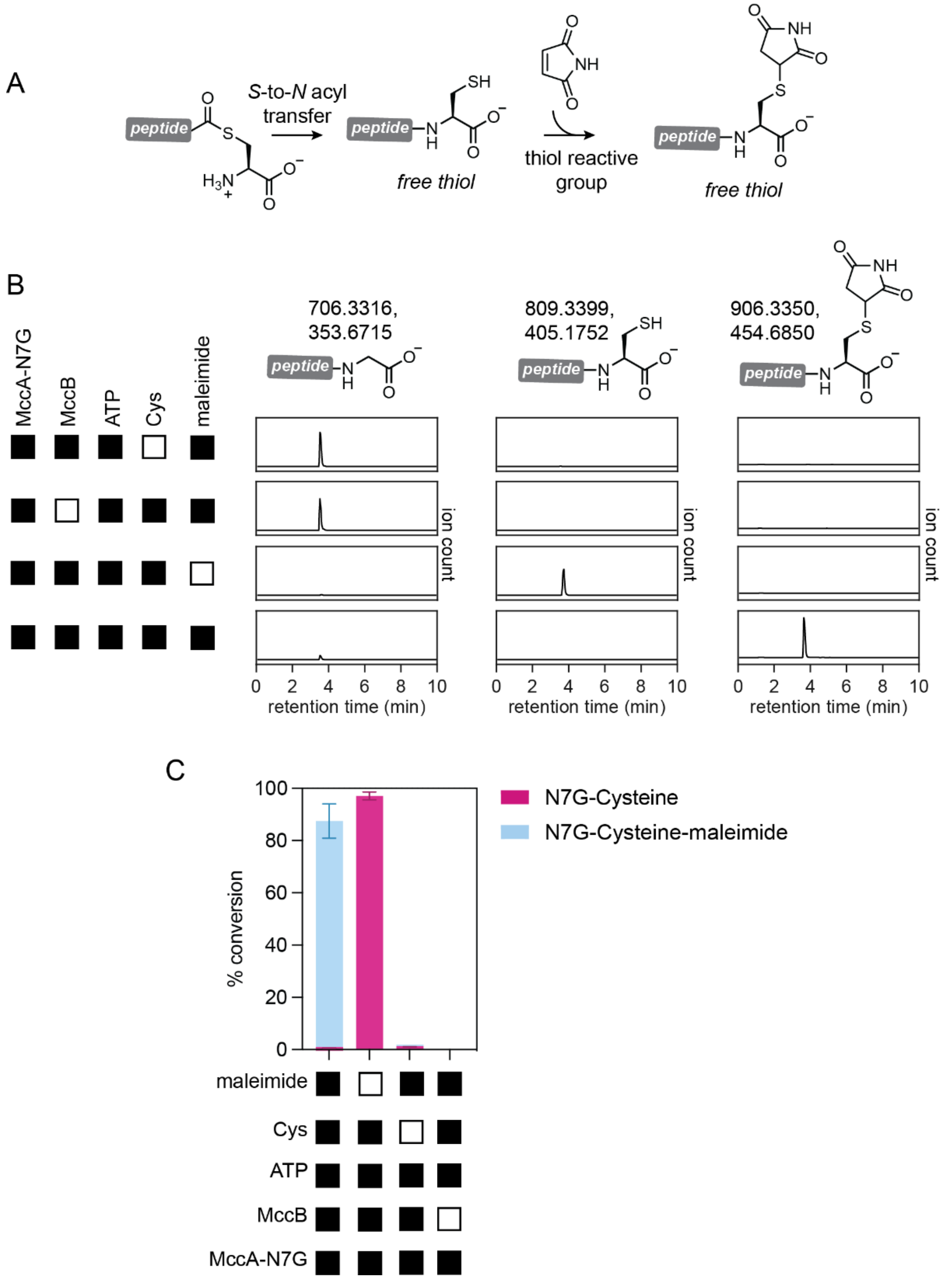

**Figure S8. Native chemical ligation (NCL) application of MccA-N7G Mesna thioester with CysTrp dipeptide.** (A) Scheme of initial Mesna thioesterification of MccA-N7G followed by transthiolation with a CysTrp dipeptide. The thioester undergoes an S-to-N acyl shift to form a stable amide bond. (B) Extracted ion chromatograms of the MccA N7G reactant, Mesna thioester, and CysTrp ligation product. The masses of both singly and doubly charged species are included. (C) HPLC-MS quantification of Mesna thioesterification and CysTrp ligation.

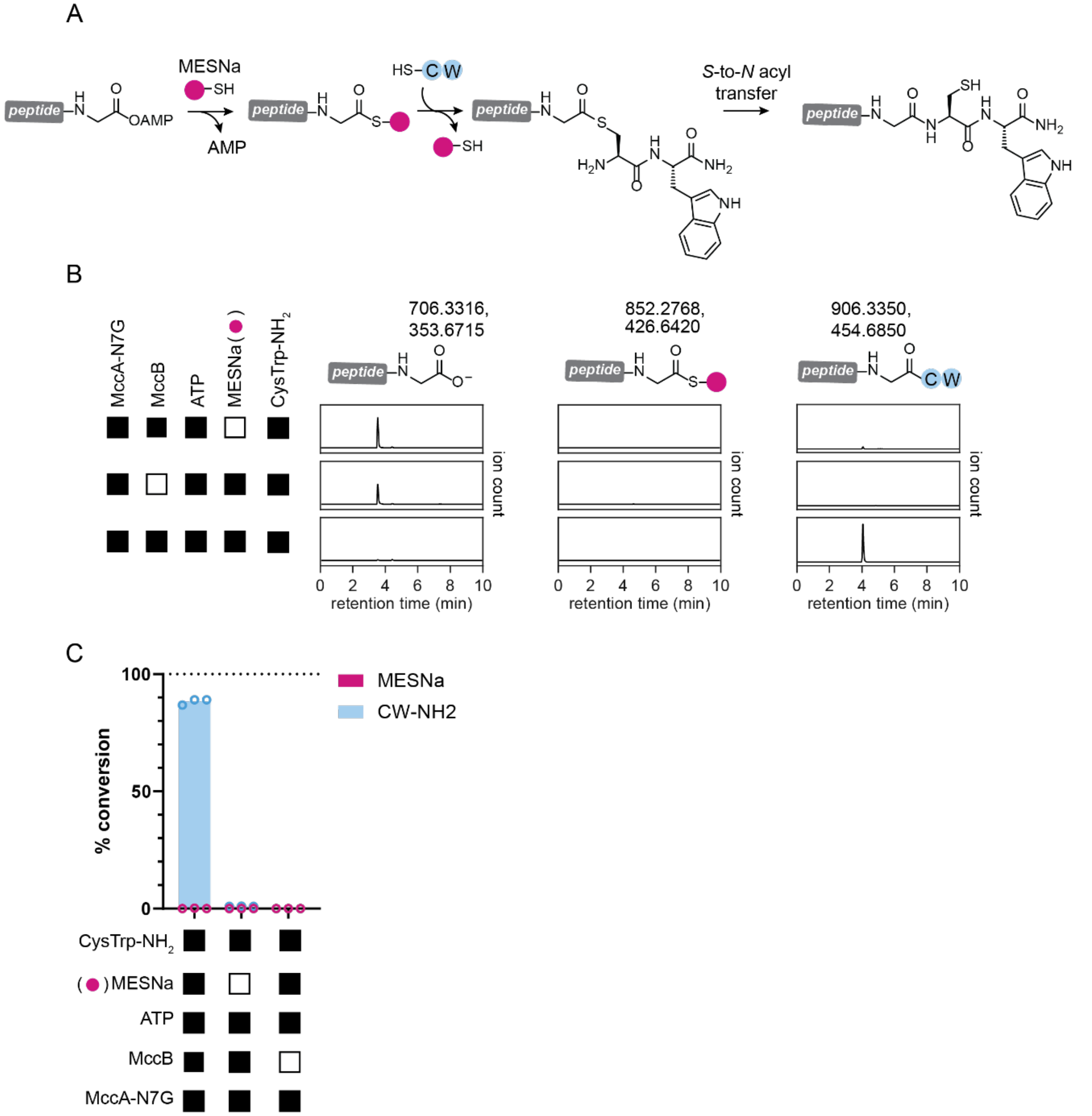

**Figure S9. Thioesterification of eGFP-TeCH using Mesna.** (A) Scheme of cysteine free eGFP-TeCH fusion thieosterification with Mesna. Reactions were performed with 5 μM MccB, 50 μM eGFP-TeCH fusion, 5 mM ATP, and the appropriate concentration of Mesna/TCEP in 75 mM Tris, pH 8.0. Reactions were quenched at selected time points by double volume addition of 0.6% TFA. (B) Representative deconvoluted traces of HPLC-MS intact protein analysis. (C) HPLC-MS quantification of Mesna thioesterification with 5 mM or 1 mM Mesna/TCEP solutions.

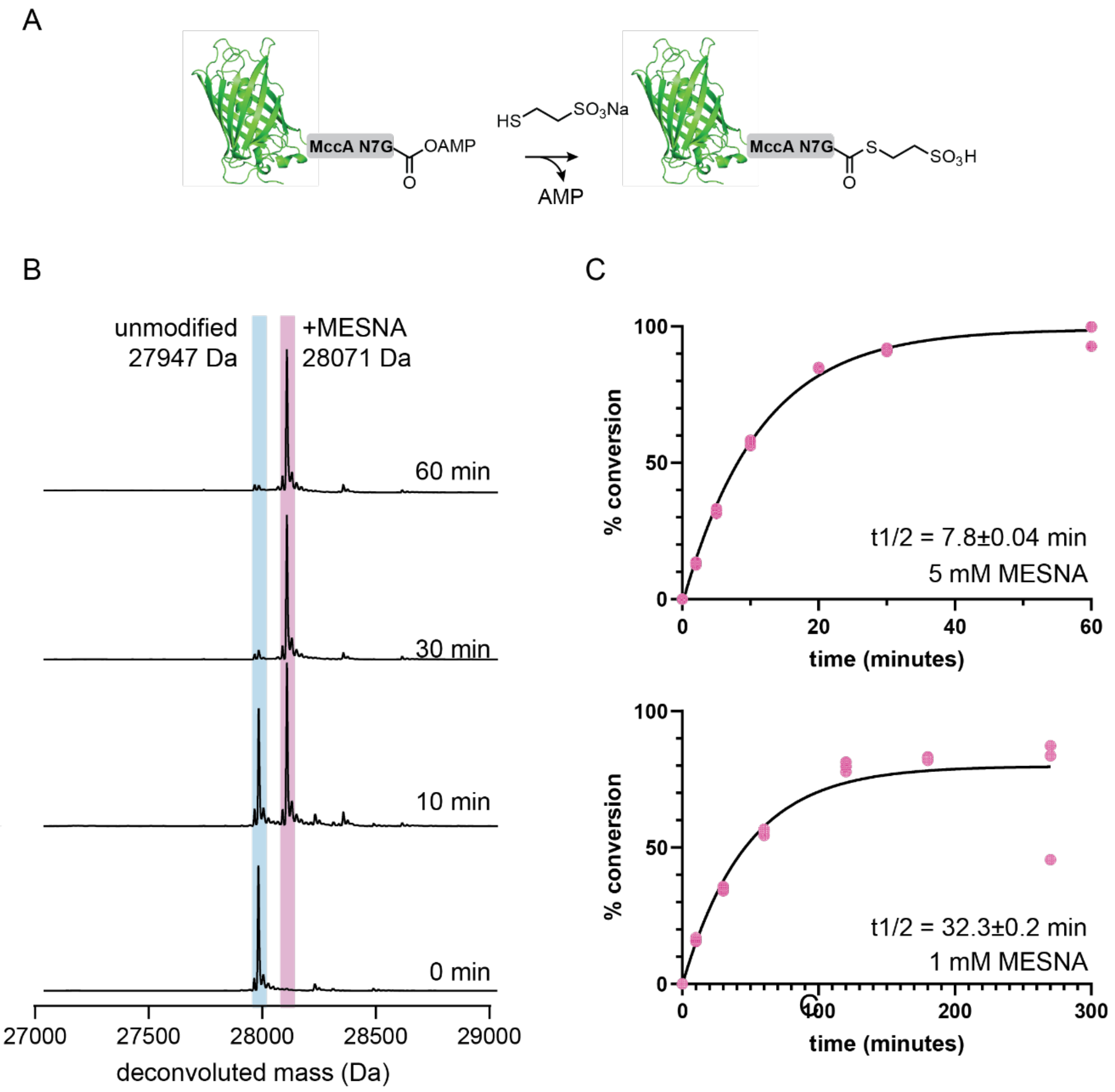

**Figure S10. C-terminal ligation of eGFP-TeCH with Cys.** (A) Scheme of cysteine free eGFP-TeCH fusion ligation with C-terminal Cys. Reactions were performed with 5 µM MccB, 50 µM eGFP-TeCH fusion, 5 mM ATP, and the appropriate concentration of Cys/TCEP in 75 mM Tris, pH 8.0. Reactions were quenched at selected time points by double volume addition of 0.6% TFA. (B) Representative deconvoluted traces of HPLC-MS intact protein analysis of eGFP-TeCH tag fusion reactions with 5 mM Cys/TCEP. (C) HPLC-MS quantification of Cys ligation with 5 mM or 1 mM Cys/TCEP solutions.

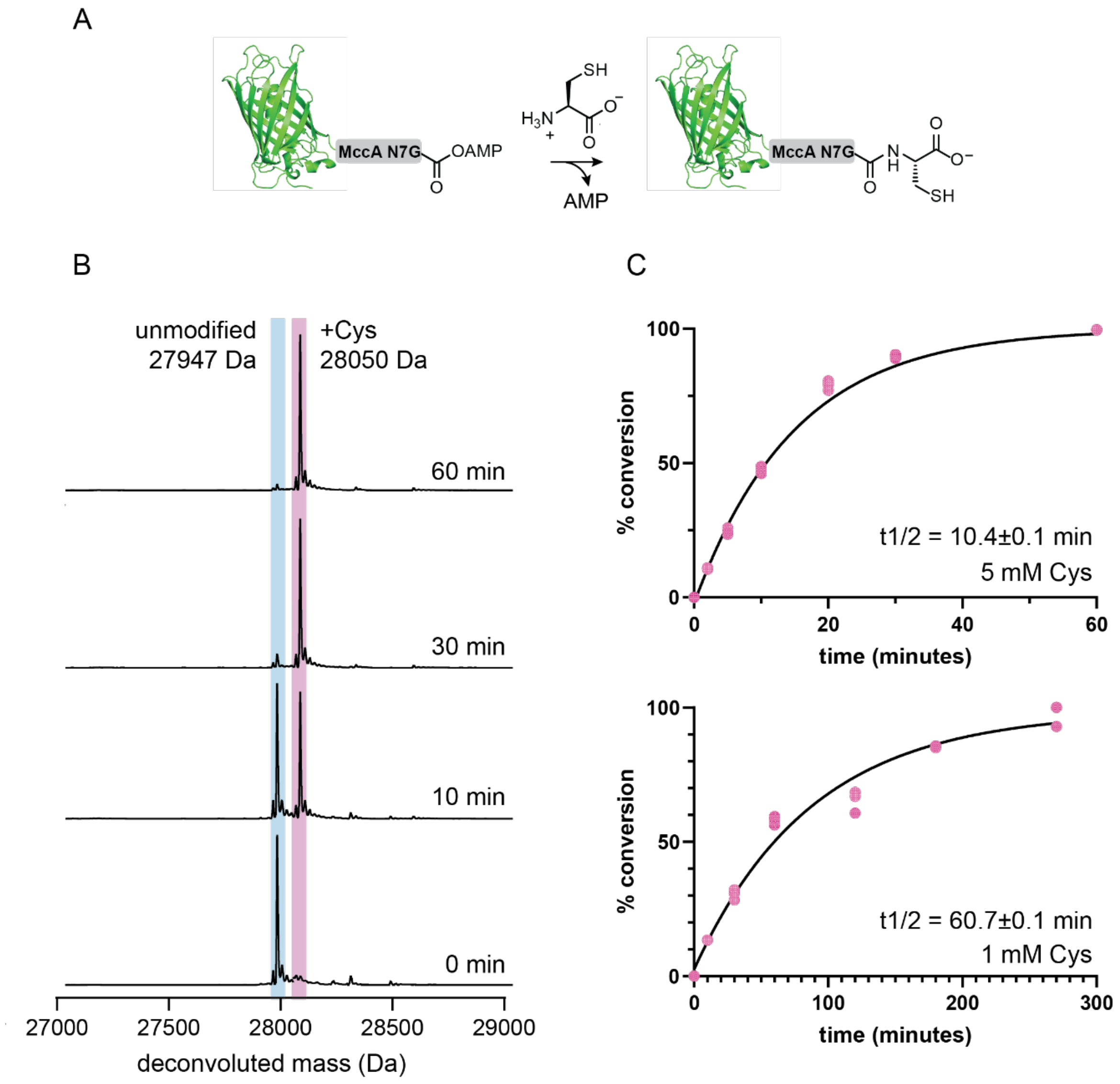

**Figure S11. Native chemical ligation (NCL) application of eGFP-TeCH Mesna thioester with CysTrp dipeptide.** (A) Scheme of cysteine free eGFP-TeCH fusion thioesterficiation followed by transthiolation/*S*-to-*N* acyl shift of CysTrp dipeptide. Reactions were performed with 5 µM MccB, 50 µM eGFP-TeCH fusion, 5 mM ATP, 5 mM Mesna/TCEP, and 5 mM CysTrp dipeptide in 75 mM Tris, pH 8.0. Reactions were quenched at selected time points by double volume addition of 0.6% TFA. (B) Representative deconvoluted traces of HPLC-MS intact protein analysis. (C) HPLC-MS quantification of Mesna thioesterification and CysTrp ligation over time.

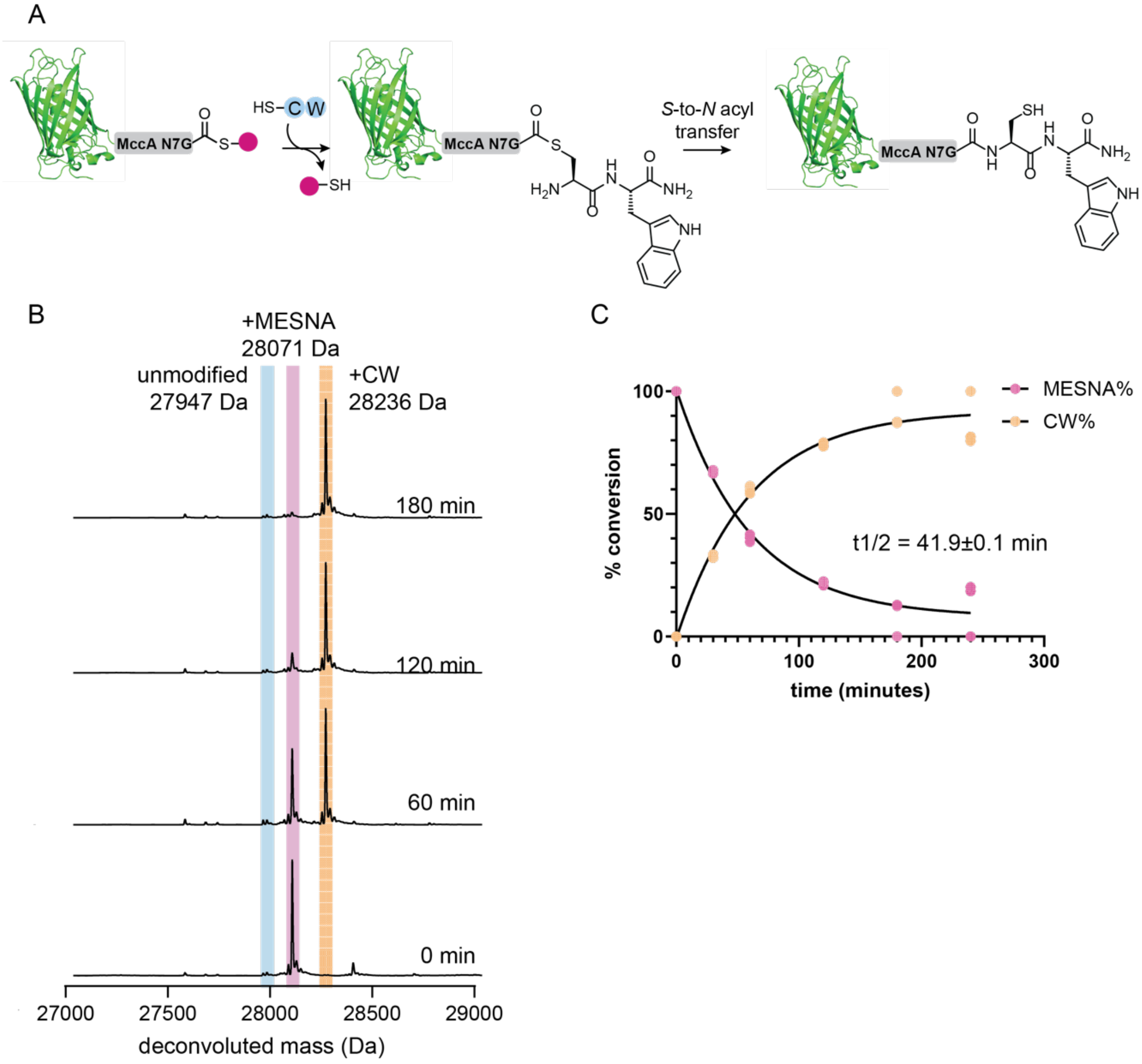

**Figure S12. Expressed protein enzyme catalyzed protein ligation (EPL) with eGFP-TeCH and AlaPhe dipeptide.** (A) Scheme of one pot reaction for eGFP-TECH thioesterficaition followed by Subtiligase-catalyzed ligation with AlaPhe dipeptide. (B) HPLC-MS quantification of AlaPhe ligation product with 5 mM AlaPhe. (C) Representative deconvoluted traces of HPLC-MS intact protein analysis. (C) HPLC-MS quantification of Mesna thioesterification and CysTrp ligation with 5 mM AlaPhe. (D) HPLC-MS quantification of AlaPhe ligation product with titrated AlaPhe concentrations. (E) Representative deconvoluted traces of HPLC-MS intact protein analysis of eGFP-TeCH expressed protein EPL with decreasing concentrations of AlaPhe.

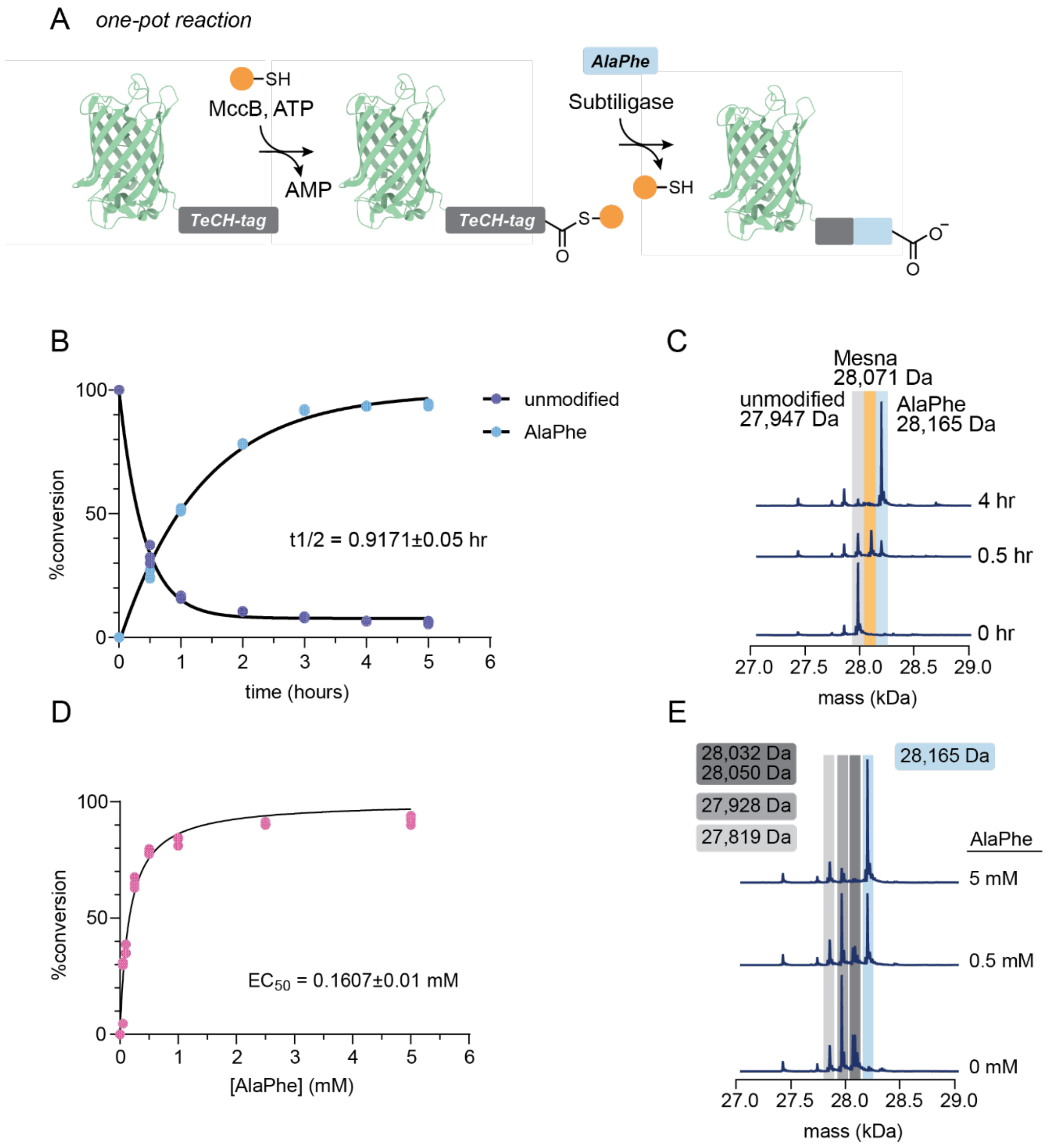

**Figure S13. Preparation of α-GFP TeCH with Cy3 conjugation.** (A) α-GFP TeCH ligation with an azidoLys containing peptide followed by SPAAC with DBCO-Cy3. Reactions were performed with 5 µM MccB, 50 µM eGFP-TeCH fusion, 5 mM ATP, 5 mM Mesna, and 5 mM azidoLys peptide in 75 mM HEPES, pH 8.0. Reactions were buffer exchanged into 75 mM HEPES three times using Zeba spin desalting columns (Thermo Scientific). Proteins were modified using SPAAC by incubating with 5 molar equivalents of DBCO Cy3 for 4 hours at room temperature, then buffer exchanging twice into 75 mM HEPES using Zeba sping desalting columns. (B) Workflow for doxycycline induction of surface eGFP expression followed by α-GFP Cy3 staining.

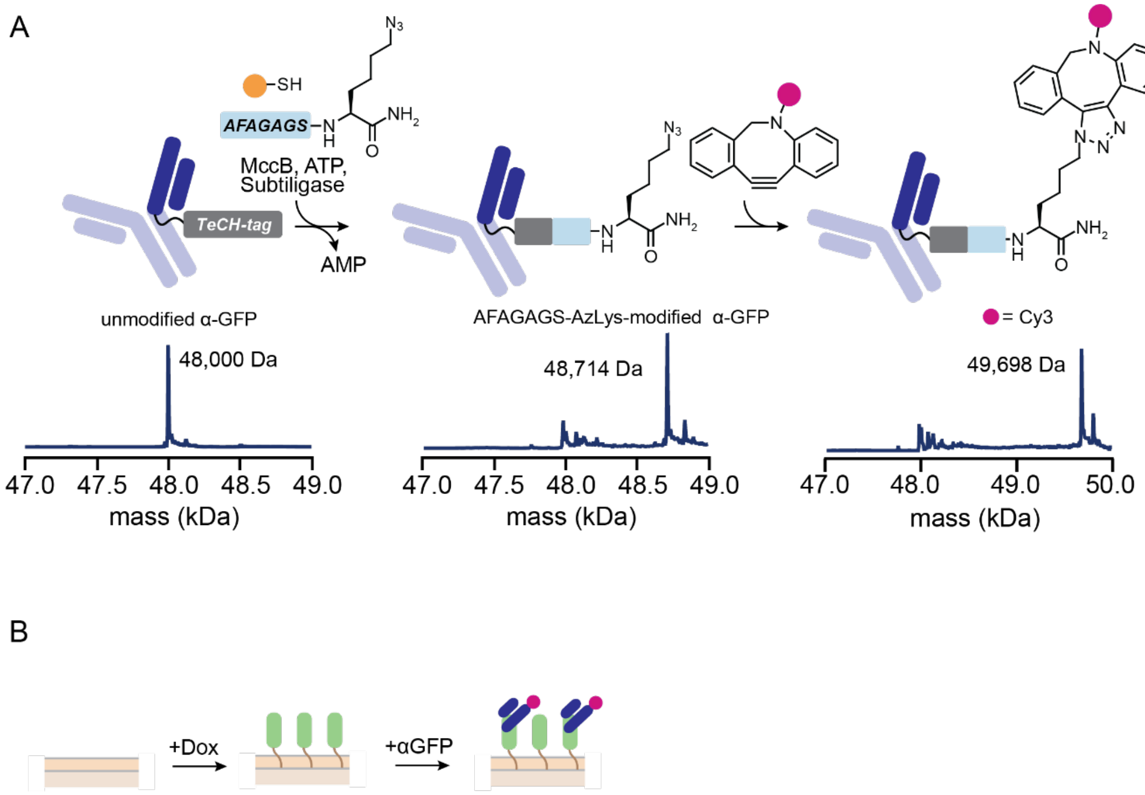

**Figure S14. Quantification of conversion of MccA positional scanning peptide library variants to *N*-AMPylated product.** Reactions contained 5 μM MccB, 250 μM MccA variant, and 5 mM ATP and were incubated for 16 h at room temperature. Percent conversion was calculated using the formula (product peak area)/(product peak area + reactant peak area)*100. (A) M1X variants. (B) R2X variants. (C) T3X variants. (D) G4X variants. (E) N5X variants. (F) A6X variants. (G) N7X variants.

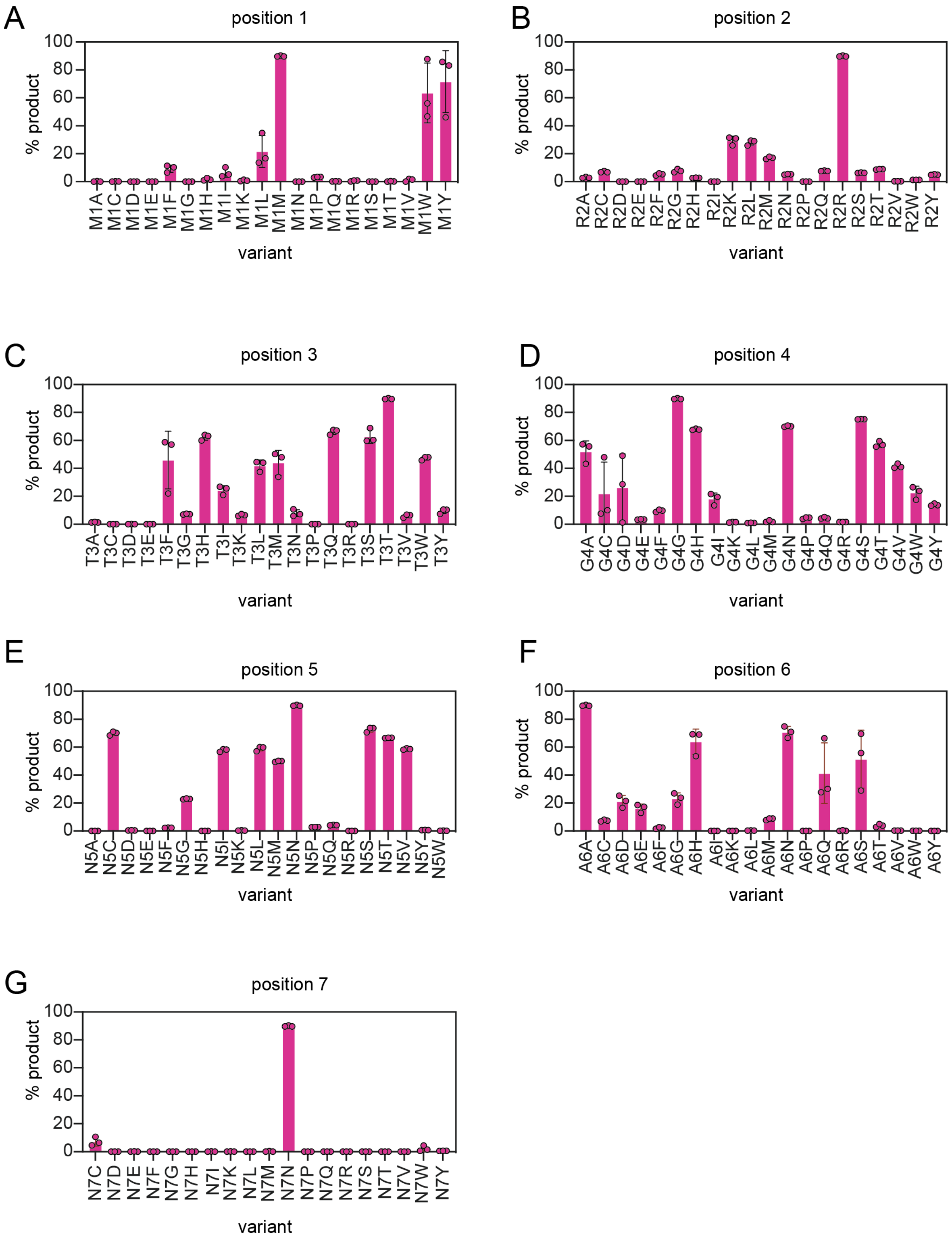

**Figure S15. Activity of *E. coli, H. pylori*, and *L. johnsonii* MccB homologs toward *E. coli, H. pylori*, and *L. johnsonii* MccAs measured with an enzyme-coupled assay for pyrophosphate release.** (A) Rate of pyrophosphate release catalyzed by MccB in presence of 250 µM of the corresponding MccA variant. Absorbance is converted to molarity using the MESG extinction coefficient. (B) Raw absorbance values used to calculate rates.

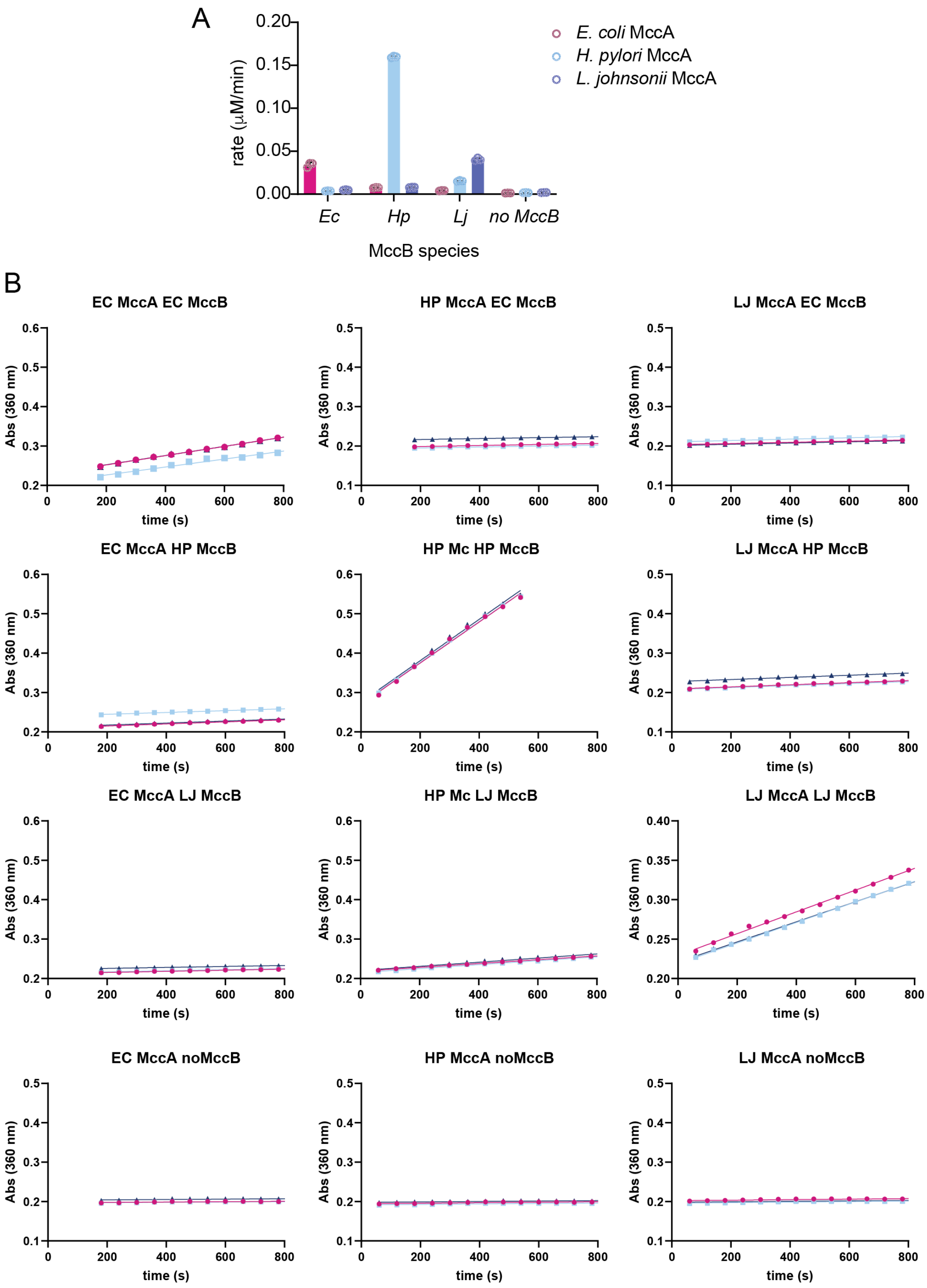

**Figure S16. Activity of *E. coli, H. pylori*, and *L. johnsonii* MccB homologs toward *E. coli, H. pylori*, and *L. johnsonii* MccA-N7Gs measured with an enzyme-coupled assay for pyrophosphate release.** (A) Rate of pyrophosphate release catalyzed by MccB in presence of 250 µM of the corresponding MccA variant. Absorbance is converted to molarity using the MESG extinction coefficient. (B) Raw absorbance values used to calculate rates.

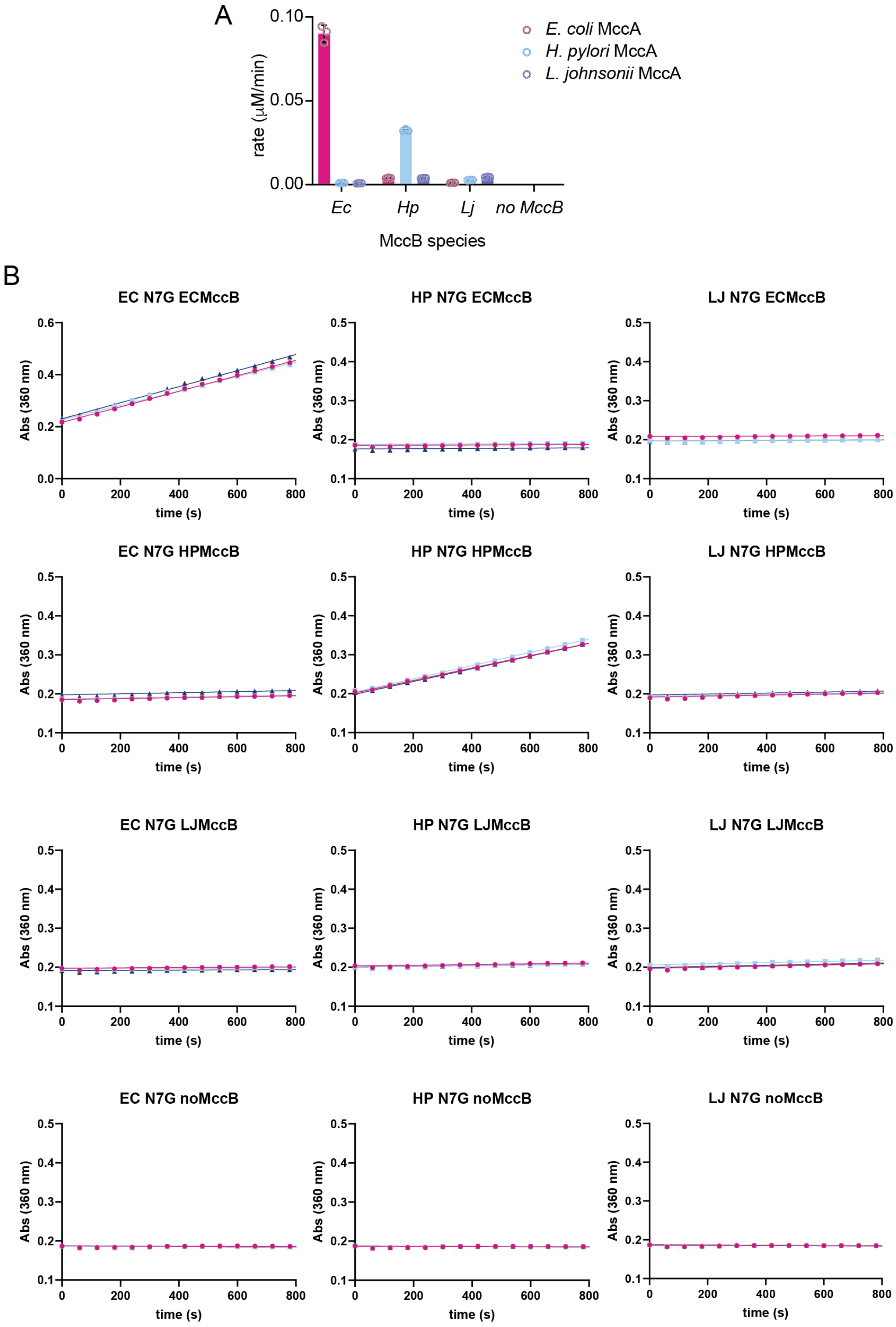

**Figure S17. C-terminal thioesterification activity of *E. coli, H. pylori*, and *L. johnsonii* MccB homologs measured by LC-TOF MS.** Reactions contained 5 μM MccB, 250 μM MccA-N7G, 100 mM Mesna, and 5 mM ATP and were incubated for 16 h at room temperature.

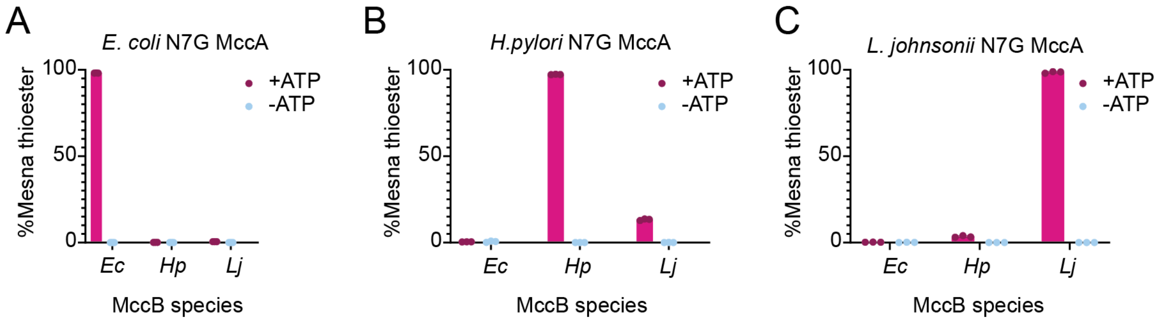

**Figure S18. C-terminal *N*-AMPylation activity of *E. coli, H. pylori*, and *L. johnsonii* MccB homologs measured by LC-TOF MS.** Reactions contained 5 μM MccB, 250 μM MccA, and 5 mM ATP and were incubated for 16 h at room temperature.

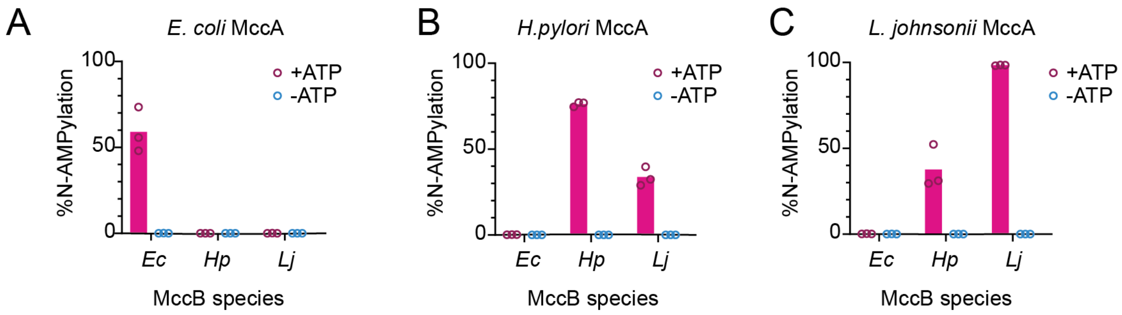

**Figure S19. Protein purification gels.** (A) Bioconjugation enzymes. (B) TeCH-tagged GFPs. (C) Panel of TeCH-tagged protein substrates.

A

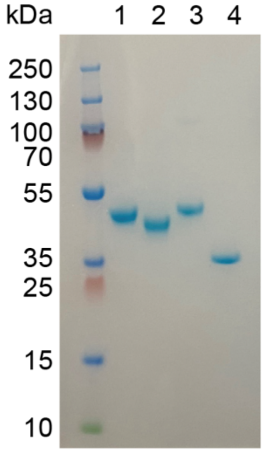


1: *E. coli* MccB, 41871 Da
2: *H. pylori* MccB, 40183 Da
3: *L. johnsonii* MccB, 42480 Da
4: subtiligase, 28545 Da

B

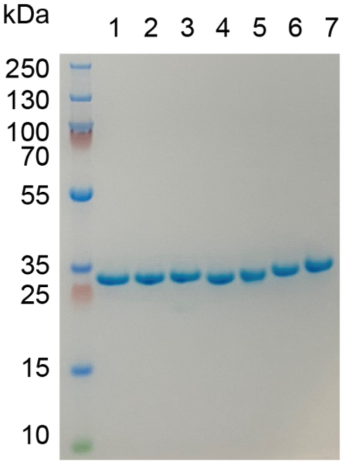


1: cf-eGFP1xTeCH, 27966 Da
2: cf-eGFP1xTeCH *H. pylori*, 28114 Da
3: 2: cf-eGFP1xTeCH *L. johnsonii*, 28132 Da
4: eGFPTeCH, 27773 Da
5: eGFP1xTeCH, 28001 Da
6: eGFP2xTeCH, 28289 Da
7: eGFP3xTeCH, 28548 Da

C

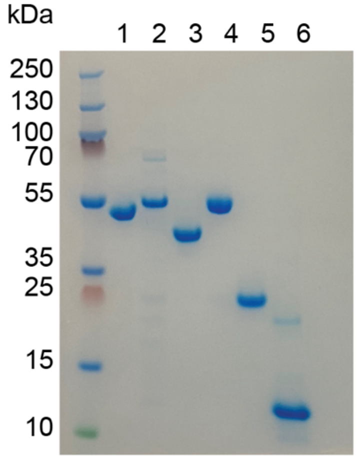


1: MBP-1xTeCH, 43599 Da
2: protein L-1xTeCH, 42578 Da
3: PTP1B-1xTeCH, 38372 Da
4: HCFab-1xTeCH, 48001 Da
5: HCFab-1xTeCH reduced
6: zEGFR-1xTeCH, 10701 Da
